# Mating system of *Ustilago esculenta* and its polymorphism

**DOI:** 10.1101/384727

**Authors:** Syun-Wun Liang, Yen-Hua Huang, Jian-Ying Chiu, Hsin-Wan Tseng, Jin-Hsing Haung, Wei-Chiang Shen

## Abstract

*Zizania latifolia* Turcz., which is mainly distributed in Asia, has had a long cultivation history as a cereal and vegetable crop. On infection with the smut fungus *Ustilago esculenta, Z. latifolia* becomes an edible vegetable, water bamboo. Two main cultivars, with a green shell and red shell, are cultivated for commercial production in Taiwan. Previous studies indicated that cultivars of *Z. latifolia* may be related to infection with *U. esculenta* isolates. However, related research is limited. The infection process of the corn smut fungus *Ustilago maydis* is coupled with sexual development and under control of the mating type locus. Thus, we aimed to use knowledge of *U. maydis* to reveal the mating system of *U. esculenta*. We collected water bamboo and isolated 145 *U. esculenta* strains from Taiwan’s major production areas. By using PCR and idiomorph screening among meiotic offspring and field isolates, we identified three idiomorphs of the mating type locus and found no sequence recombination between them. Whole-genome sequencing (Illumina and Pacbio) suggested that the mating system of *U. esculenta* was bipolar. Mating type locus 1 (*MAT-1*) was 555,862 bp, and contained 44% repeated sequences. Sequence comparison revealed that *U. esculenta MAT-1* shared better conservation with the sex chromosome of *U. maydis* than *U. hordei*. These results can be utilized to further explore the genomic diversity of *U. esculenta* isolates and their application for water bamboo breeding.

## INTRODUCTION

Water bamboo is one of the most popular vegetables in Asia. When the wild rice *Zizania latifolia* Turcz. is infected by the smut fungus *Ustilago esculenta*, tumor-like tissue is produced in the basal stem of plants. This gall tumor is known as “water bamboo”, ‘Kal-peh-soon’ or ‘Jiaobai’. The red shell water bamboo cultivar, a single-season plant, and green shell cultivar, a multi-season plant, are two main cultivars in Taiwan. However, with LED technology, the green shell cultivar is now harvested year-round. The major difference in the two cultivars is that the red shell cultivar grows with red spots on the inner layer of the leaf sheath and the other lacks these spots.

The cultivar variation is caused by the different characteristics of *U. esculenta* strains (Hung *et al*. 2001). Because *U. esculenta* invades host plants, its hyphae can be observed in the rhizome and basal stem of its host plant, including nodes, buds and shoots, which have been explored thoroughly by fluorescence microscopy (Jose *et al*. 2016). Meanwhile, auxin and cytokinin are accumulated after *U. esculenta* infection, followed by the induction of galls on the host plant.

The concentration of auxin shows a gradient variation in different stages of gall tumor (Chan and Thrower 1980; Chung and Tzeng 2004; Wang *et al*. 2017). During hyphae invasion, both intercellular and intracellular processes exist. With intercellular invasion, hyphae penetrate the plant cell and digest its interior contents. Several hyphae then gather in this space, followed by the production of teliospores from the ends of the hyphae (Zhang *et al*. 2012).

*U. esculenta* is a smut fungi belonging to the Basidiomycota. It has several common characteristics with other smut fungi, such as dimorphism of cell morphology, production of powder-like teliospores and invading the host plant as a sexual hypha (Martinez-Espinoza 1993; Kües *et al*. 2011; Ye *et al*. 2017). On the basis of the origin of gall characteristics, *U. esculenta* strains are designated as teliospore (T) and mycelia-teliospore (M-T) strains. ‘Baishin’, ‘Huashin’ and ‘Heishin’ are three types of galls found in Taiwan. Heishin usually produces the T strain, producing abundant teliospore sori in the early gall formation stage, whereas Huashin or Baishin produces the M-T strain, with few or no teliospores produced in the later gall formation stage (Yang and Leu 1978; Zhang *et al*. 2017). Baishin has high economical value as compared with Huashin. However, Heishin is discarded once found in the field. Therefore, *U. esculenta* is forced to maintain in the mycelial state and fails to complete sexual reproduction inside the vegetatively propagated crop host. *U. esculenta* lacks several essential virulence factors and has had relatively slow evolution under many years of artificial selection (Ye *et al*. 2017). The genome of the diploid mycelium in *U. esculenta* was sequenced and analyzed.

The mating system in several fungi has been linked to the infection; examples are *Ustilago maydis* (Bakkeren *et al*. 2008) and *Cryptococcus neoformans* (Nielsen and Heitman 2007). In the broad study of fungus, the determinant genes are conserved and have been reviewed: mating factor (*mfa*), pheromone receptor (*pra*), and heterodimeric homeodomain transcription factors (HD1 and HD2) (bE and *bW* genes in *U. maydis*) (Brefort *et al*. 2009). When two opposite mating-type cells meet, one pheromone is recognized by another pheromone receptor. *Mfa* encodes a precursor pheromone peptide, which is sensed and received by a pheromone receptor. The Pra protein then triggers the MAPK module and Prf1 protein to activate the expression of the bE/bW heterodimer complex (encoded by *bE* and *bW* genes). The subsequent pathogenic progress begins, including development of the conjugation tube and penetration. (Anderson *et al*. 1999; Brefort *et al*. 2009; Kües *et al*. 2011)

Sexual behavior in the fungi is known as homothallism or heterothallism. Heterothallism is further classified as bipolar and tetrapolar based on the absence of genetic linkage of a and b gene regions. A gene region mainly includes *pra* and *mfa* genes, whereas b gene region contains *bE* and *bW* genes. Several smut fungi identified by mating type include *U. maydis* (2 a and 25 b gene complex) and *Sporisorium reilianum* (3 a and 5 b gene complex), which are tetrapolar, and *Ustilago hordei*, which is bipolar (Stakman and Christensen 1927; Schirawski *et al*. 2005; Kües *et al*. 2011). *U. hordei* carries the largest mating-type loci in smut fungi: *MAT-1* and *MAT-2*, which are 500 and 430 kb, respectively (Bakkeren and Kronstad 1994; Lee *et al*. 1999). As well, 50% of long terminal repeats and transposable elements appear on *MAT-1* (Horns *et al*. 2012).

The origin of the mating system is controversial. Genome comparison of these regions inferred that *U. maydis* and *U. hordei* both evolved from *S. reilianum*, so the evolution of the mating system was from tetrapolar to bipolar (Bakkeren *et al*. 2006; Laurie *et al*. 2012). The *U. esculenta* mating system is considered tetrapolar (Yang and Leu 1978) and carries three gene complexes and three b gene complexes (Ye *et al*. 2017). However, we lack solid evidence for this.

Every year, farmers face huge production loss because of the formation of teliospores in water bamboo for unknown reasons. Without understanding the process of infection, solving this problem is difficult. This study examined the mating type locus of *U. esculenta* and three other smut fungi. To further understand the *U. esculenta* mating process and its possible application for reducing agricultural production loss, we aimed to reveal the complete mating type locus of *U. esculenta* and its system.

## MATERIALS AND METHODS

### Isolation and preservation of *U. esculenta*

Two main cultivars, green shell and red shell, of *Zizania latifolia*-infected samples were collected from the commercial fields and an agricultural research institute in Taiwan. The red shell cultivar is mainly distributed in northern Taiwan, and the green shell cultivar is predominantly cultivated in central Taiwan. *U. esculenta* was isolated from the galls of infected plants by micromanipulation or tissue isolation. Micromanipulation was used to isolate strains from galls with teliospore sori. In brief, teliospores were collected from infected tissues and suspended in sterile water. Spores were spread onto potato dextrose agar (PDA) containing 50 μg/ml chloramphenicol to induce germination at 28°. About 18 to 20 hr post-incubation, haploid meiotic progeny were picked by micromanipulator (ECLIPSE 50i, Nikon) when most promycelia contained four sporidia. Tissue isolation was used to isolate strains from galls lacking black sori. Leaf sheaths were first removed and intact galls were collected and cut open under laminar flow. Ten pieces of internal tissue were aseptically excised, placed on PDA medium containing 50 μg/ml chloramphenicol, and incubated at 28° for 1 to 2 weeks. Sporidial colony produced from the sections were then re-suspended, diluted and spread on PDA medium. Finally, haploid sporidial strain was obtained from a single colony. Strains were preserved by culturing strains at 28° in potato dextrose broth (PDB) plus 1% sorbose for 72 hr. Culture was then mixed with an equal volume of cryoprotective liquid (20% glycerol, 10% lactose), kept at -20° for 30 min and then stored at -80°.

### Nuclear and septal staining

Nuclear and septal staining were performed by using 4’, 6-diamidino-2-phenylindole (DAPI; Sigma) and calcofluor white (CFW; fluorescent brightener 28, Sigma). *U. esculenta* cells were first fixed in fixation buffer (3.7% formaldehyde, 0.1 M phosphate buffer, 0.2% Triton) for 30 min, then rinsed with sterile distilled water. Samples were stained with CFW (10 μg/ml) for 5 min and washed with distilled water, then stained with DAPI (0.8 μg/ml) for 30 min and finally de-stained with distilled water. Stained samples were examined by fluorescent microscopy (Olympus BX41) with a filter set (Olympus, U-MWU2 BP330~385).

### Mating assay

Haploid *U. esculenta* strains subjected to mating assay were cultured on PDA medium at 28° for 3 days. Haploid yeast strains were individually re-suspended in sterile water and mixed in pairs with an equal amount. Then, a 5-μl cell mixture was spotted onto PDA medium containing 1% charcoal. After 10 to 14 days, if two tested strains were compatible for mating, mating hyphae developed around the edge of the colony. Alternatively, for mating assay conducted on GMM medium (Shimizu and Keller 2001), mating filaments were formed and visible within 3 to 5 days. Photos of mating colony were taken by camera (Coolpix P300, Nikon).

### DNA extraction

To extract *U. esculenta* genomic DNA, strains were freshly grown in PDB with agitation at 28° for 60 hr. Cells were harvested by centrifugation and lyophilized. Cell materials of about 0.1 g were re-suspended in 500 μl of 65° pre-warmed CTAB buffer (2% CTAB, 1.4 M NaCl, 20 mM EDTA, 100 mM Tris pH 8, 2% PVP-40) and 3 μl mercaptoethanol was added. Samples were incubated at 65° for 30 min and mixed every 10 min by inverting the tubes. After 30 min, 500 μl phenol/chloroform (1:1 volume ratio) was added. Samples were gently mixed and then spun at 15600 g for 15 min. After centrifugation, the upper aqueous phase was transferred to a new tube and an equal volume of chloroform was added. Samples were gently mixed and spun at 15600 g for 5 min. The upper aqueous phase was transferred to a new tube, and 0.7 times the volume of isopropanol was added. Samples were inverted for 10 times, kept on ice for 10 min and underwent centrifugation at 15600 g for 5 min. Supernatants were discarded, and pellets were washed by adding 500 μl of 75% ethanol. Samples were spun at 15600 g for 5 min and supernatants were carefully removed. DNA pellets were air-dried, and 100 μl of sterile distilled water was added to re-suspend the DNA. DNA concentration was measured by spectrophotometry (NanoDrop 1000, Thermo Fisher Scientific, USA) and DNA samples were stored at -20°.

For PCR screening, DNA template was prepared by the fast preparation of fungal DNA (FPFD) method (Liu *et al*. 2011). First, a small quantity of yeast cells was suspended in 100 μl extraction buffer (15 mM Na_2_CO_3_, 35 mM NaHCO_3_, 2% PVP40, 0.2% BSA, 0.05% Tween20), then incubated at 95° for 15 min. The tube was immediately kept on ice and spun down by centrifugation. Then, 3 μl supernatant was used as template DNA for PCR reactions.

### Identification of *U. esculenta* mating type-related genes by PCR

To identify the mating type genes of *U. esculenta*, several PCR approaches were used. Sequences for evolutionally conserved genes related to mating type locus and flanking regions, including *bE, pra, lba1, panC, c1d1* and *Nat1*, from related smut fungi were aligned. Conserved regions were identified, and specific or degenerate PCR primers were designed for PCR amplification as described (Albert and Schenck 1996). PCR fragments with expected sizes were purified and underwent TA cloning (pGEM-T Easy, Promega). Transformants were screened and verified by sequencing. To amplify the complete A/B mating type locus, long-range PCR was conducted with primers designed on the flanking genes of the mating type locus. PCR conditions followed that suggested in the product manual (Q5 Hot Start High-Fidelity DNA Polymerase, New England Biolabs; TaKaRa LA Taq DNA Polymerase, TaKaRa Bio). Six *U. esculenta* isolates, 12JK1RB1-A1 (a1b1), 12JK1RB1-A2 (a3b3), 12SB1RB1-B4 (a2b2), 13PJ1GB1-D3 (a1b1), 13PJ1GB1-D4 (a2b2), and 13PJ3GB1-E1 (a3b3), were selected to determine the sequences of mating type locus.

### Screening of *U. esculenta* mating type by multiplex PCR

Three sets of idiomorph specific primers for mating type A and B gene clusters were designed to screen the mating type of *U. esculenta* isolates. Multiplex PCR reaction was performed as follows: denaturation at 94° for 3 min, 35 cycles of denaturation at 94° for 45 sec, annealing at 55° for 30 sec, and extension at 72° for 25 sec, and final extension at 72° for 7 min.

### Illumina sequencing of *U. esculenta* genome

To prepare high-quality genomic DNA for next-generation sequencing, a modified CTAB method was used (Winnepenninckx *et al*. 1993). Briefly, *U. esculenta* strains were first grown in PDB for 60 hr and cells were collected by centrifugation and lyophilized, then suspended in 800 μl of 60° pre-warmed CTAB buffer (2% CTAB, 1.4 M NaCl, 20 mM EDTA, 100 mM pH 8 Tris, 2% PVP-40) plus 0.2% mercaptoethanol and 0.1 mg/ml proteinase K. Samples were incubated at 60° for 1 hr and gently inverted every 20 min. After incubation, 800 μl chloroform/isoamyl alcohol (24:1) was added and gently mixed for 2 min. Samples were then centrifuged at 14000 g for 10 min at 4°. The upper aqueous phase was transferred to a new tube, and 1 μl RNase (100 μg/μl) was added and kept at 37° for 90 min, then 600 μl isopropanol was added, mixed gently, and samples were kept at room temperature overnight to allow DNA precipitation. The next day, samples were centrifuged at 14000 g for 15 min at 4°. Supernatants were discarded and pellets were washed by adding 800 μl of 75% ethanol. Samples were spun for 5 min at 15600 g, and supernatants were removed completely and pellets were air-dried. Finally, 100 μl sterile distilled water was added to re-suspend DNA pellet. DNA samples were quantified by Qubit (Invitrogen) and the concentration was adjusted to 10 ng/μl. For DNA shearing, Covaris S2 (Covaris, MA, USA) was used to break DNA into 200-bp fragments; fragments 200 to 700 bp were selected by using Ampure XP beads (Beckman Coulter Genomics, CA, USA).

Library construction involved the Illumina TruSeq DNA kit. First, the ends of size-selected fragments were repaired, and poly A nucleotides were added to 3’ ends. Poly T complemented specific adaptors were linked to both ends of fragments and 10 cycles of PCR amplification were conducted. The distribution of fragment size was confirmed by BioAnalyzer before next-generation sequencing (Agilent Technologies, CA, USA). Pair-ended sequencing with 250-bp reads was conducted with Illumina MiSeq (Illumina Inc., CA, USA) at the NGS Core (Center for Systems Biology, National Taiwan University).

The cluster density of sequencing was 1105 K/mm^2^ and size of 12JK1RB1-A1 and 12JK1RB1-A2 was 5.5 and 4.6 Gb. Three assembly programs were used: CLC bio, SOAPdenovo (Short Oligonucleotide Analysis Package) and velvet. Performance was assessed by read mapping rate and number of orthologous proteins between *U. esculenta* and *U. maydis*. The assembled results with CLC bio were better than with other methods and were used for gene annotation. The average base coverages were 230 X and 194 X for 12JK1RB1-A1 and 12JK1RB1-A2.

### Long-read PacBio sequencing

The method was modified from the Gentra kit DNA extraction procedure. Briefly, 3-day-old *U. esculenta* strains were first incubated in 5 ml PDB for 24 hr and then transferred to a 125-ml flask and grown for another 34 hr (no longer than 36 hr). Cells were then counted to approximate 2 x 10^8^ cells. First, samples were suspended in 300 μl cell suspension solution and 60 mg VinoTaste (Novozymes), followed by reaction at 37° for 1 hr. After that, 300 μl cell lysis solution plus 1% SDS was added for incubation at 50° for another 30 min. Then, 100 μl protein precipitation solution was added, followed by vigorously vortexing for 20 sec, then centrifugation for 3 min at 16,000 g, then DNA was pelleted by inverting the tubes 50 times with the addition of 300 μl isopropanol. Pure DNA was acquired by washing with 70% Ethanol, 5 to 10 min air drying and re-suspended with 100 μl DNA hydration solution. Finally, RNA was removed with 1.5 μl RNase A solution and incubated at 37° for 60 min for enzyme reaction, then transferred to 65° water bath for 60-min incubation.

Genomic DNA was sheared by using a Covaris g-TUBE followed by purification via binding to pre-washed AMPure PB beads (Part no. PB100-265-900). After end-repair, the blunt adapters were ligated and underwent exonuclease incubation to remove all un-ligated adapters and DNA. The final “SMRT bells” were annealed with primers and bound to the proprietary polymerase by using the PacBio DNA/Polymerase Binding Kit P6 v2 (Part no. PB100-372-700) to form the “Binding Complex”. After dilution, the library was loaded onto the instrument with the DNA Sequencing Kit 4.0 v2 (Part no. PB100-612-400) and 4 SMRT Cells 8Pac for sequencing. A primary filtering analysis was performed with the RS instrument, and the secondary analysis involved using the SMRT analysis pipeline v2.3.0.

### Bioinformatics analysis

Protein coding genes of *U. esculenta* were predicted by using GeneMark-ES (Borodovsky and Lomsadze 2011) to scan the assembled contigs, then the predicted genes were annotated in a similarity-based manner. NCBI-BLASTP (Altschul *et al*. 1990) was used to search for homologous genes in UniRef clusters (Suzek *et al*. 2015). When finding significantly similar hits to a predicted gene (*i.e.*, with e-value < 10^-4^, low complexity filtering by using SEG program), the annotation information was retrieved from the best hit; otherwise, HMMer and Pfam HMM models were used to predict the protein domains in the encoded amino-acid sequences of the genes remaining unannotated in the similarity-based annotation step (Durbin *et al*. 1998; Finn *et al*. 2016).

Repeat and transposable elements were predicted by using RepeatModeler 1.0.8 plus the RMBlast search tool (Smit and Hubley; Smit *et al.*) and TransposonPSI (Haas). Part of the genome comparison analysis was carried out by using MUMmer 3.23 (Kurtz *et al*. 2004) and diagrammed by using Easyfig 2.2 (Sullivan *et al*. 2011). Comparative analysis of the UE genome with 10 other fully-sequenced fungal genomes, including *Aspergillus nidulans*, *Cryptococcus neoformans, Magnaporthe oryzae, Neurospora crassa, Phytophthora infestans*, *Puccinia graminis, Saccharomyces cerevisiae, Sporisorium reilianum, Ustilago maydis* and *Ustilago hordei*, involved using the Ensembl Compara pipelines, with inter-species syntenic regions and phylogenetic trees of all protein-coding genes inferred by using the Lastz-net pipeline (Kent *et al*. 2003) and the GeneTrees pipeline (Vilella *et al*. 2009), respectively. All the results of genome annotation and comparative analysis were integrated into a locally maintained Ensembl genome database and browser system for further data mining and visualization.

### Inoculation experiment

For inoculation experiments, *U. esculenta* isolates were first cultured on PDA medium at 28° for 5 to 7 days. Uninfected *Z. latifolia* plants collected from the field were used for inoculation. Collected plants were initially washed clean with tap water. Unwanted leaves, leaf sheaths, and roots were removed, and 15 to 20 cm long of basal stems containing young buds were saved. Basal stem tissues were further washed with distilled water, kept in a plastic box containing perlite and distilled water, then grown in a controlled growth chamber with the conditions of 24°, 75% relative humidity, 8 hr daylight with light intensity 75 μmol s^-1^ m^-2^ for 3 to 5 days to maintain viability. After incubation, the buds showing vigorous viability were selected for inoculation. Outer layers of leaf sheath surrounding the buds were removed and two tiny holes pierced into one bud tissue were created by using an insect-pinning pin. Yeast cells of compatible *U. esculenta* isolates were mixed in equal amount and applied to wounded sites on buds. The inoculated samples were placed back into a perlite box and tightly covered with a plastic bag to retain humidity. The plastic bag was dislodged 3 days later, and inoculated samples were maintained in a controlled growth chamber at 24°, 75% relative humidity, 8-hr daylight with light intensity 75 μmol s^-1^ m^-2^. After 2 weeks, surviving buds were excised, transferred to soil pots and grown in a greenhouse. Evaluation of successful inoculation rate was conducted 4 to 6 weeks post-inoculation by examining whether swelling of the basal stem occurred.

### Data availability

All data necessary for confirming the conclusion of the article are present in the article, figures and tables. Sequencing data used in this research was deposited in the NCBI (data will be submitted to NCBI later). Figure S1 presents the reads coverage of genome of *MAT-1*. Figure S2 illustrates the syntenic region of *MAT* loci between *U. esculenta* and *U. hordei*. Table S1 and S2 are *U. esculenta* strains isolated from the fields. Table S3 presents the primer sets used in this study.

## RESULTS

### Field plant collection and isolation of *U. esculenta*

Two field cultivars, red and green shells, were collected from northern Taiwan (Jinshin) and central Taiwan (Puli), respectively (Figure 1A). According to the presence of teliospores, these collections were classified into three groups: Baishin, Huashin and Heishin. Baishin refers to a snowy white gall without teliospores (Figure 1B, C). Heishin is known by a huge number of teliospores scattered inside the gall, whereas Huashin has fewer teliospores. Furthermore, Heishin is smaller and grows faster than Huashin during the harvest time (Figure 1D).

**Figure 1.**
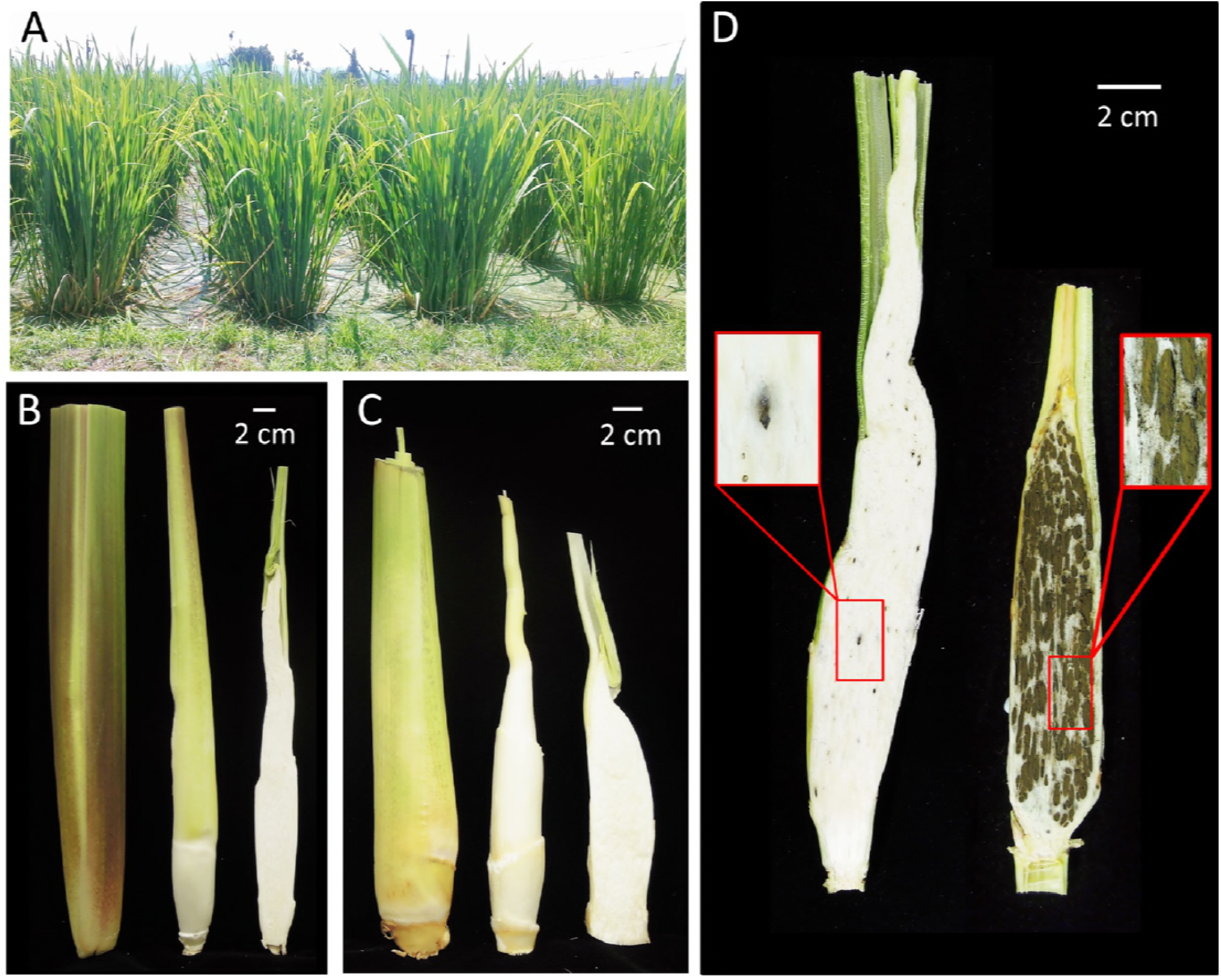
Water bamboo plants in the commercial field and features of each edible gall in different Taiwan varieties. (A) Red shell variety of water bamboo was cultivated in Sanchi, Taiwan. (B) Features of the red shell variety. From left to right: appearance of edible gall, gall without leaf sheath, longitudinal section of gall. (C) Features of the green shell variety. From left to right: appearance of edible gall, gall without leaves sheath, longitudinal section of gall. (D) Swelling galls with black teliospore sori. Mature galls of red shell variety collected from Sanchi show different levels of sori formation; left: sori scattered in the swelling tissue, or huashin, right: gall full of sori, or heishin.

*U. esculenta* strains were isolated by micromanipulation or directly from plant tissue. With the former method, 59 strains were separated from Heishin tissue, including 44 strains from 11 sets of four meiotic progenies and 15 strains from 7 incomplete sets. Strains from Baishin and Huashin were directly isolated from plant tissue: 9 from Baishin tissue and 77 from Huashin tissue (Table S1, S2, S3).

### Life cycle of *U. esculenta*

Most smut fungi share a similar life cycle pattern; however, *U. esculenta* is slightly different from others because of the artificial vegetative propagation. For more than 100 years, farmers have tried to avoid the development of teliospores in the paddy field. Thus, most *U. esculenta* strains in the wild remain in the hyphae state. Under natural circumstances, as for other smut fungi, *U. esculenta* infects host plants as filaments and produces tumor-like tissue and teliospores (Figure 2A). Teliospores germinated and produced 2 septa, then evolved to 5 or 6 septa in promycelium while 4 meiotic yeast-like sporidia were produced. The fourth sporidia usually budded from the teliospore directly. Likewise, sporidia, which were unicellular or multicellular, also performed an *in vitro* asexual cycle. When opposite sporidia were conjugated and formed sexual filaments, we found three characteristics in the mating filaments by using DAPI and CFW stains. In young filaments, the front end region included living cells and dead cells, which were present as empty-cytoplasm sections. In the late stage, several dikaryotic cells appeared at the tip region, and the monokaryotic yeast cells budded from the growth tip and the edge of some septa (Figure 2E-2H).

**Figure 2.**
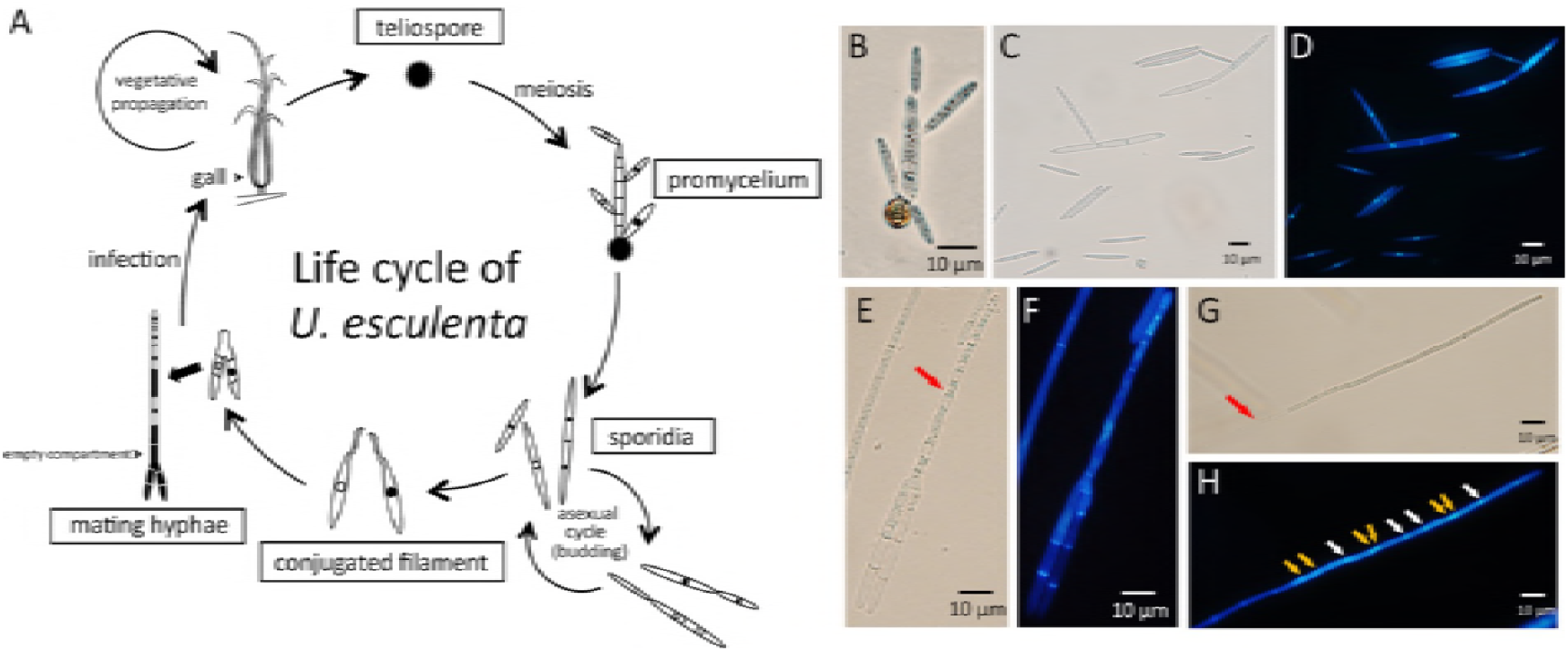
Life cycle and morphological features of *U. esculenta*. (A) The life cycle of *U. esculenta* is associated with infection of *Zizania latifolia* Turcz. In the asexual cycle, *U. esculenta* grows as yeast/sporidia and reproduces asexually by budding. In the sexual cycle, two compatible sporidial cells produce conjugated filaments to fuse and generate mating hyphae. Budding yeast cells at the front parts and empty cell compartments are usually observed. Mating filaments penetrate into host plants and develop swelling galls at the basal stem, which was stimulated to grow by the disturbance of phytohormone homeostasis. Teliospore sori are then produced in the interior of mature galls. Teliospores are diploid, which gave rise to promycelium, followed by meiosis and produce four haploid yeast cells with two different mating types. (B) Teliospores collected from infected plants were induced to germinate on PDA medium. Several septa and four sporidia were observed after 20 hr. (C) *U. esculenta* sporidia existed as single or multiple yeast formed cells. (D) DAPI staining of sporidia in sample (C) revealed a single nucleus in each cell. (E) Two compatible haploid sporidia conjugated to produce mating hypha (red arrow). (F) DAPI and CFW staining of sample (E) show the positions of nuclei and septa. (G) Mating hyphae of *U. esculenta* were characterized by empty cell compartment (red arrow). The right top was the direction of hyphal growth. (H) DAPI and CFW staining of sample (G) show both mononuclear and binuclear cells at the front end. Monokaryons and dikaryons are indicated with white and yellow arrows, respectively.

### *U. esculenta* features heterothallism

*U. esculenta* is known as a heterothallic fungi. To reveal the mating type system, we used mating assay by inter/intra-mating assay of 6 sets of teliospore-isolated strains, including 3 sets from green shell cultivars (13PJ1GB1, 13PJ3GB1, 13PS2GB1) and 3 sets from red shell cultivars (2 from 12SB1RB1 and 1 from 12JK1RB1). For the intra-mating results, 4 of 6 crossings showed furry colonies (Figure 3A), so two different mating types were present in the 4 meiotic progenies. For the inter-mating results, both 8 and 12 compatible crossings were observed among the 16 crossings (Figure 3B, C). The former result featured two different mating type loci in 8 strains, and the later result more than 2 mating types (Table 1).

**Figure 3.**
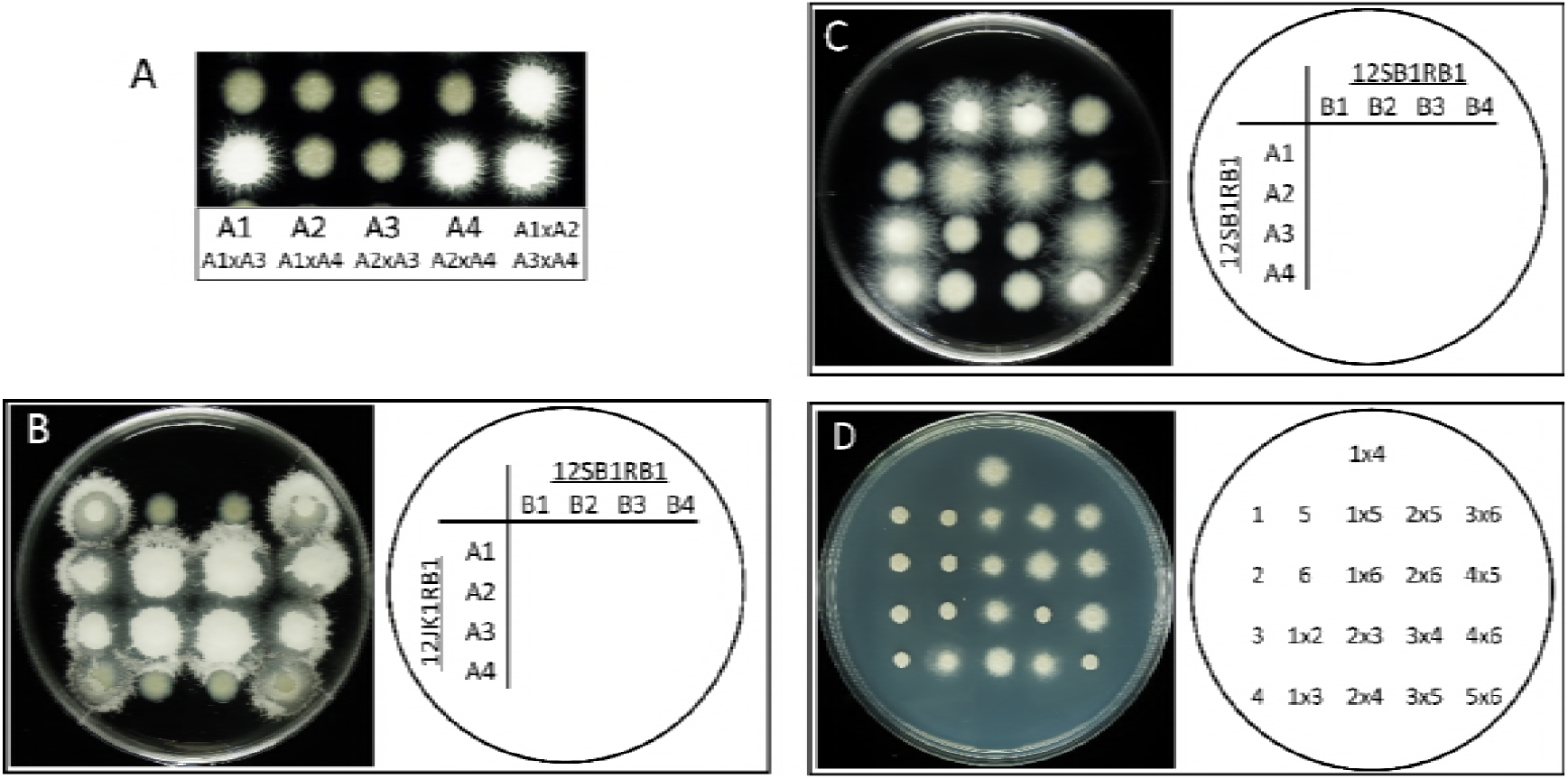
Mating assay of *U. esculenta* isolates derived from the meiotic progenies of teliospores. (A) Mating assay of the 4 meiotic progenies of teliospore 12JK1RB1_A. Four of 6 pairs showed compatible results according to the production of mating filaments. (B) Mating result of 8 strains germinated from two teliospores: 12JK1RB1_A and 12SB1RB1_B. Twelve pairs showed compatible mating. (C) Same process but different results with different strains. Strains were produced from teliospores 12SB1RB1_A and 12SB1RB1_B. Only 8 pairs showed compatible mating. (D) Mating assay of 6 *U. esculenta* isolates whose mating type were determined. Compatible and incompatible mating pairs were confirmed (1, 12JK1RB1-A1; 2, 13PJ1GB1-A3; 3, 12SB1RB1-B4; 4, 13PJ1GB1-A4; 5. 12JK1RB1-A2; 6, 13PJ3GB1-A1). Mating assay performed on PDA medium containing 1% charcoal, except for (D) performed on GMM medium.

**Table 1.**
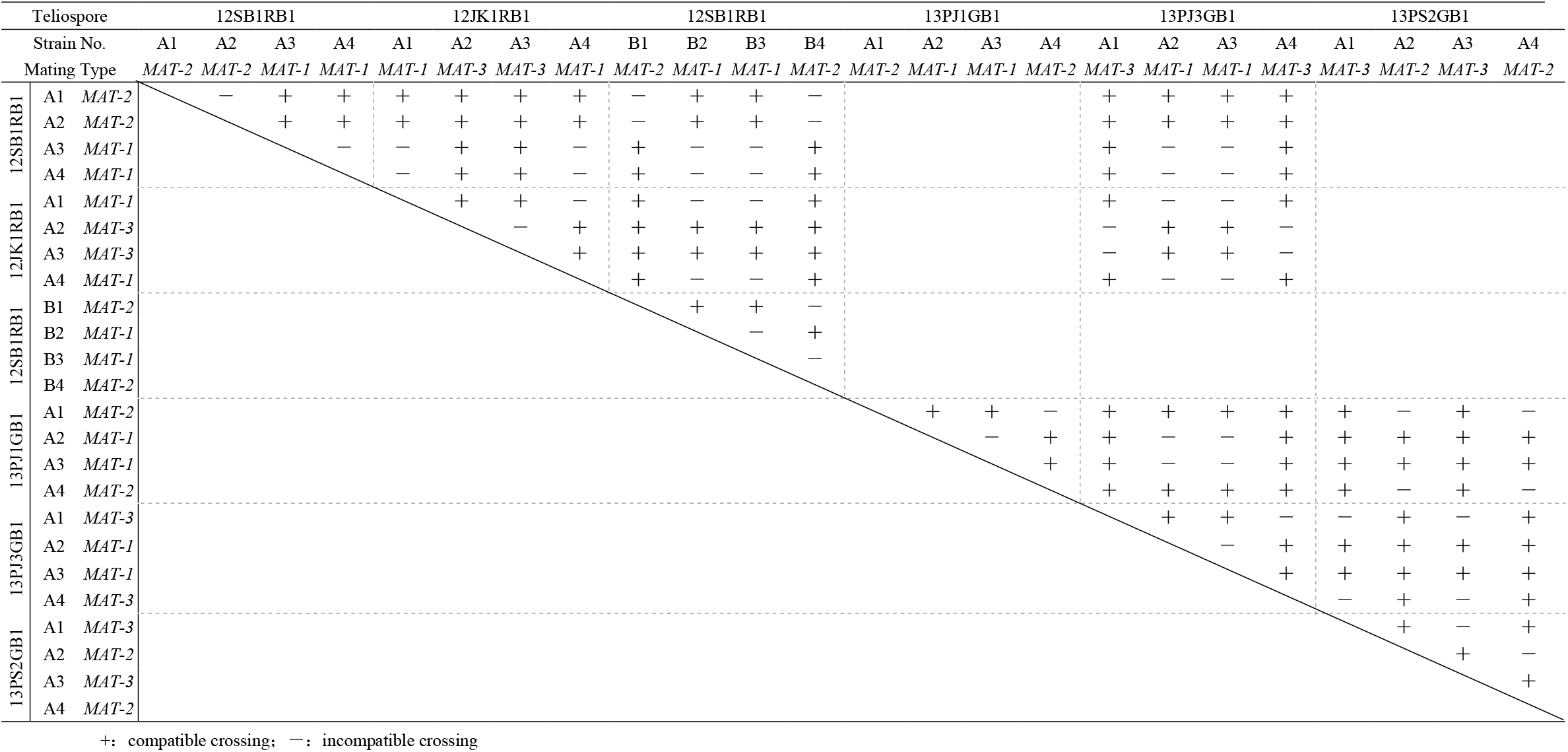
Mating assay of inter- and intra-crossing of 6 sets of teliospore-derived strains

### *U. esculenta* has 3 idiomorphs of mating type

We used the PCR primer sets for *U. scitaminea bE* gene (Albert and Schenck 1996), then designed degenerate PCR to amplify the flanking genes of A and B gene complexes. The flanking genes *c1d1, nat1, lba1* and *panC* were amplified. To obtain the complete sequence of mating type, whole-genome sequencing of 12JK1RB1-A1 (*MAT-1*) and 12JK1RB1-A2 (*MAT-3*) involved using Illumina Hi-seq.

Sequences of the non-coding region between *bE* and *bW* genes and the partial sequence of *bW* gene were highly diverse. By comparing this region among 23 field-collected strains, we found 3 different lengths of amplicons: 2,028, 2,130 and 2,141 bp, which were partial sequences of *MAT-1, MAT-2* and *MAT-3*. The primer set used was wc1529 and wc1531 (Table S4).

The full sequences of A and B gene complexes were obtained by PCR cloning or whole-genome sequencing (primer sets: Table S4). The length of three A gene complexes was 6,455, 8,129 and 7,156 bp, respectively, with B gene complexes 7,433, 7,325 and 14,165 bp respectively.

### Differences in *MAT* locus within *U. esculenta* strains and other species

A gene complex in *U. esculenta* includes several genes: pheromone receptor (*pra*), two pheromones (*mfa*) and *lga/rga* and is flanked by a left-border protein (*lba*) and right-border protein (*rba*) (Figure 4). The B gene complex, which was flanked by proposed nuclear regulator (*c1d1*) and N-terminal acetyltransferase (*Nat1*) genes, mainly contained b West (*bW*) and b East (*bE*) genes (Figure 5). Three a/b gene complexes share high synteny with other smut fungi, including *U. maydis, U. hordei*, and *S. reilianum*.

**Figure 4.**
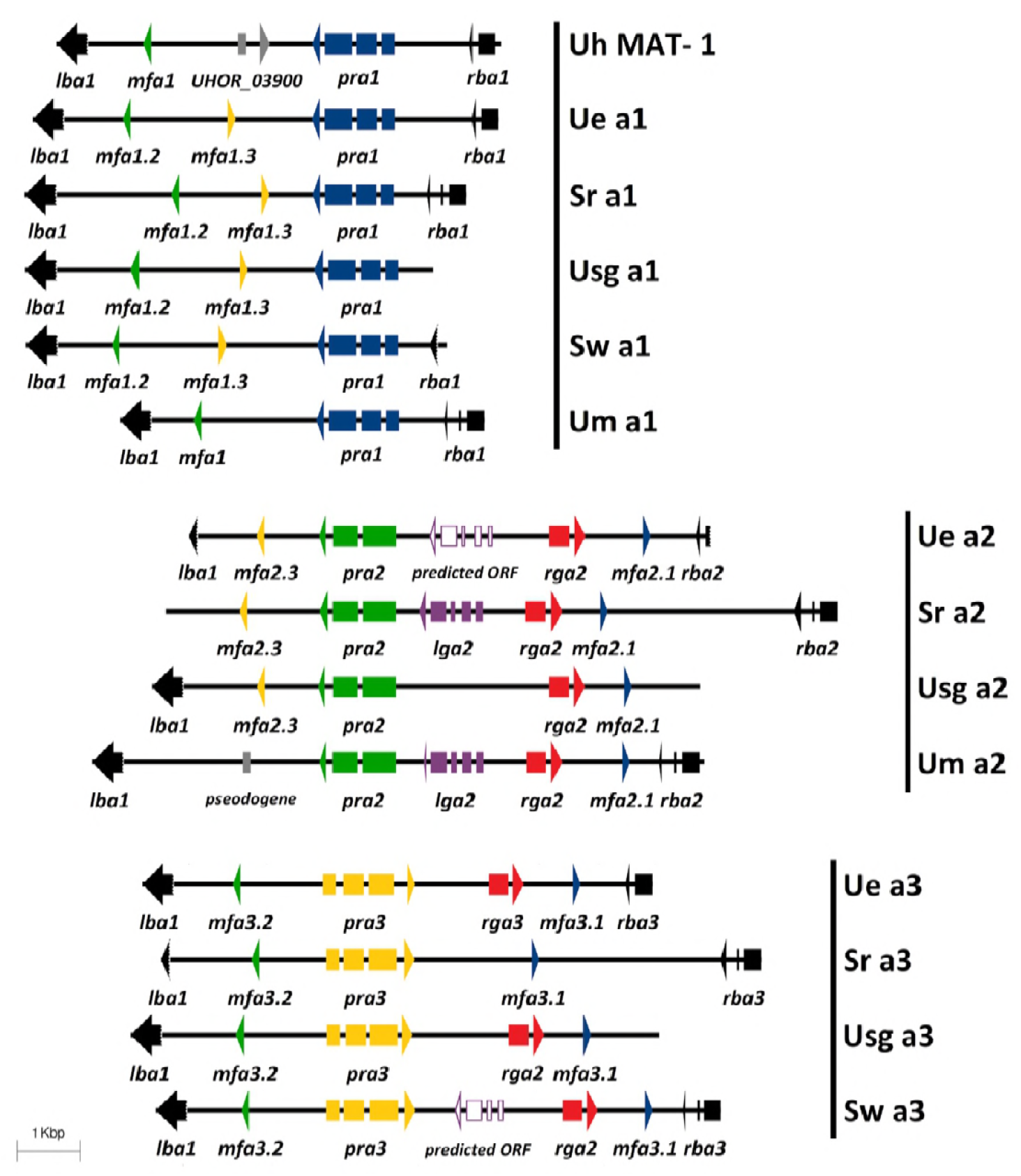
Genetic structure of mating type A region in *U. esculenta* (12JK1RB1-A1, 12SB1RB1-B4, 12JK1RB1-A2) and related smut fungi. Three clusters of A-region genes were identified among *U. esculenta* isolates in Taiwan and showed conservation with related smut fungi. Several conserved genes, including *pra* (pheromone receptor), *mfa* (mating factor/pheromone), and *lga/rga* (mitochondria inheritance-related genes), were identified and were flanked by conserved *lba* and *rba* genes. Both Ue a2 and Sw a3 have one predicted gene, annotated as a predicted open reading frame, which showed only partial similarity to other *lga* genes. (Uh, *U. hordei*; Ue, *U. esculenta*; Um, *U. maydis*; Sr, *Sporisorium reilianum*; Sw, *S. walker*; Usg, *Ustanciosporium gigantosporum*)

**Figure 5.**
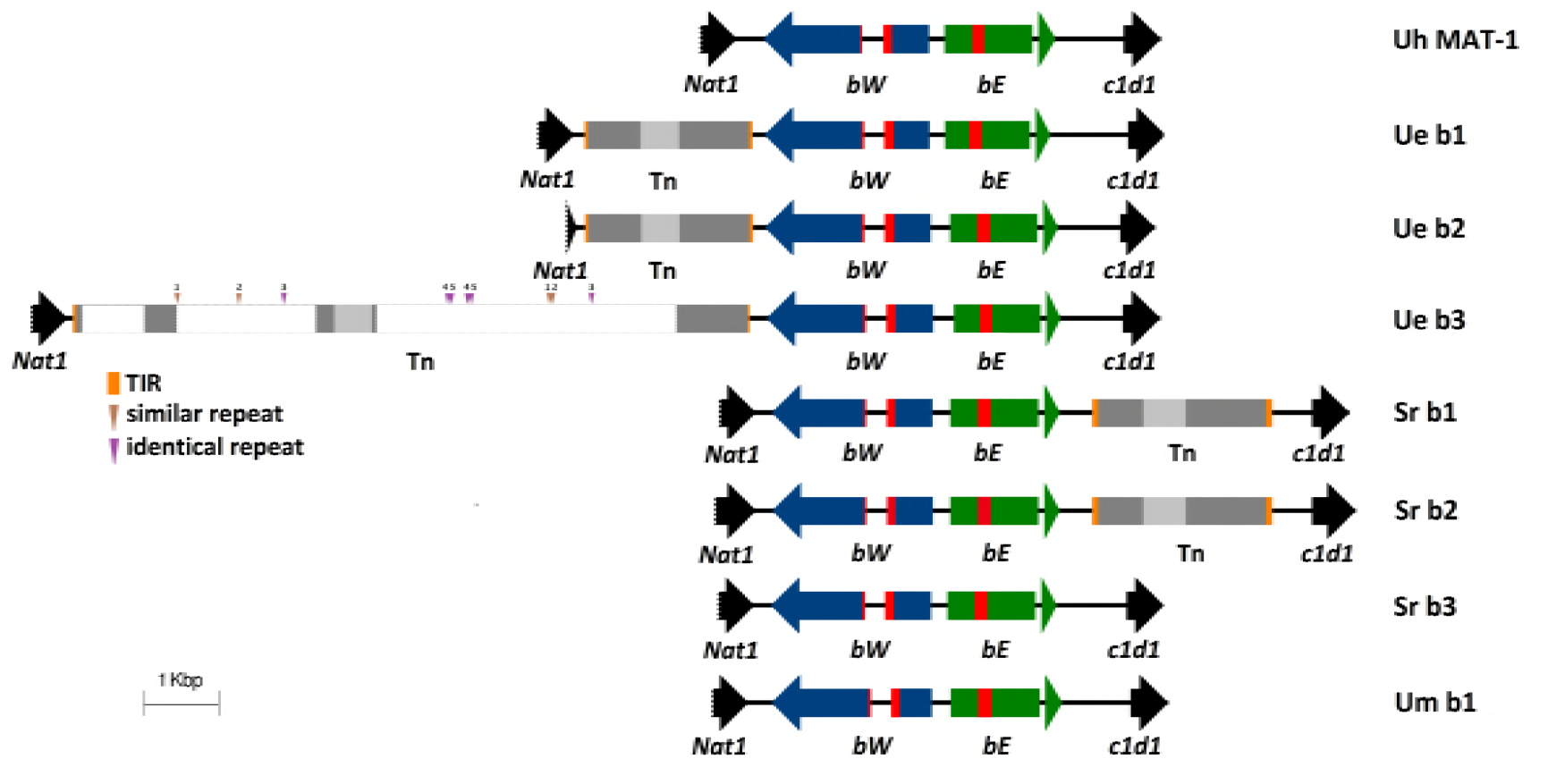
Genetic structure of mating type B region in *U. esculenta* (12JK1RB1-A1, 13PJ1GB1-A4, 12JK1RB1-A2) and related smut fungi. bWest (bW; blue arrow) and bEast (*bE*; green arrow) genes contained a homeodomain region (red block) and were flanked by the *nat1* and *c1d1* genes (black arrows). Three clusters of mating type B-related genes were identified among *U. esculenta* isolates in Taiwan. Except for common *bE* and *bW* genes, transposons were also located in this region, comprising erminal inverted repeat (TIR) (orange block) and DDE domains (light gray). Ue b3 had a large size and contained 5 different tandem repeats (arrow heads). Brown arrow indicates repeats with several single nucleotide polymorphisms and purple arrow indicates identical repeat. Identical number on the arrow indicates the same pair. The dashed white box in the b3 transposon represents the inserted region as compared to b1 and b2 transposons. *S. relianum* b1 and b2 loci carry a similar DNA transposon as well. (Uh, *U. hordei;* Ue, *U. esculenta;* Um, *U. maydis;* Sr, *Sporisorium reilianum*)

The pheromone-receptor (P/R) system of *U. esculenta* involved three Pra proteins and six Mfa proteins. Phylogenetic trees (Figure 6B, C) revealed that both Mfa and Pra proteins are divided into 3 clades. The similarity of Mfa and Pra proteins in each clade was up to 90% (76-90%) and 86% (76-86%). However, the similarity of *U. maydis* Mfa2 to other proteins in the same clade was merely 56% to 65% (Figure 6B). Also, *mfa1.3* and *mfa2.3* showed a slight difference at the N-terminus but might be identical after processing (Figure 6A).

**Figure 6.**
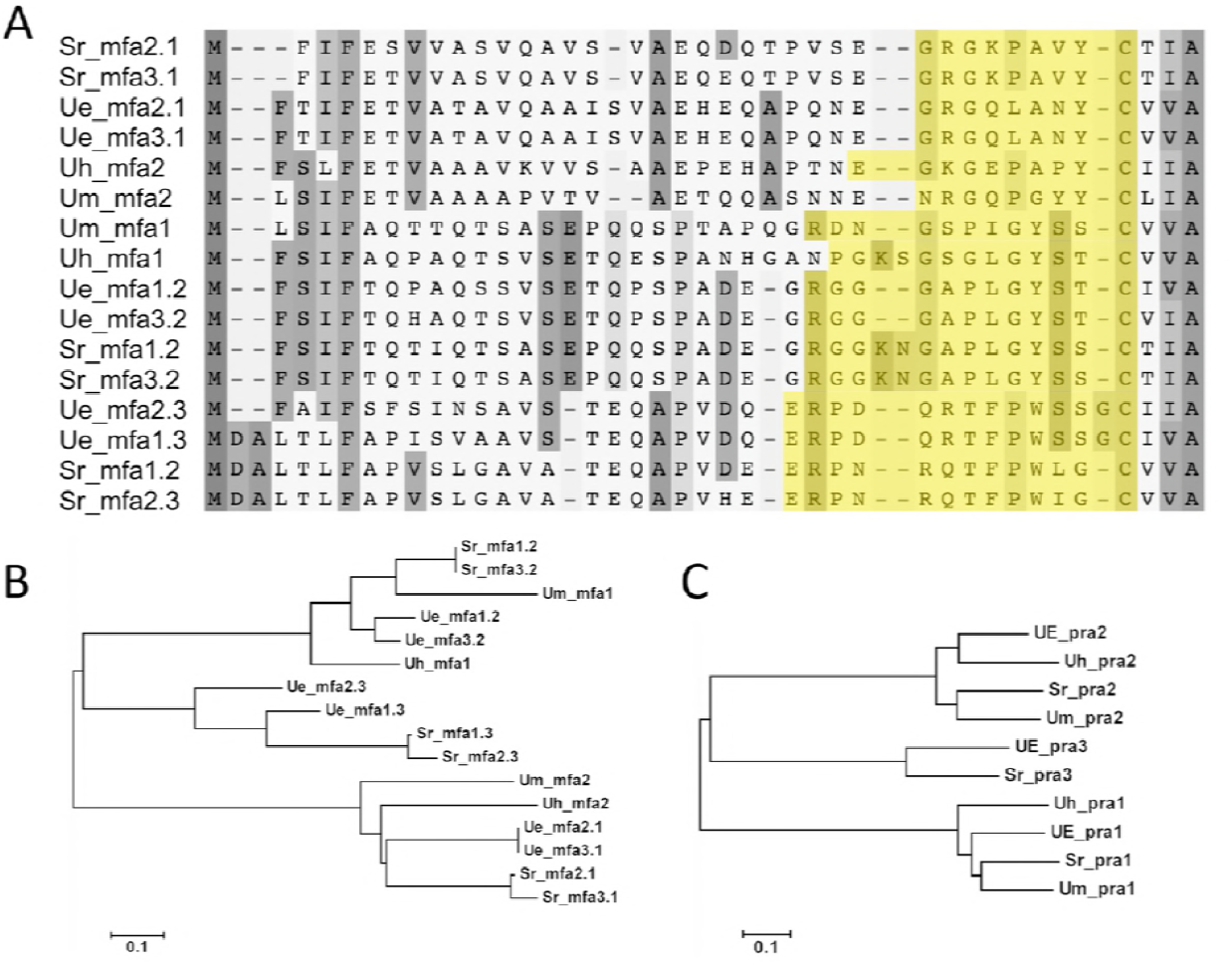
The alignments and phylogenetic analysis of *mfa* and *pra* genes. (A) Mfa protein sequence showed a conserved region of the CAAX motif and the sequence in yellow was mature pheromone protein. The protein weight matrix for alignment was point accepted mutation (PAM) and the gap penalty of both pairwise and multiple alignment was 5. (B) Phylogenetic analysis of mfa protein. (C) Phylogenetic analysis of pra protein. Both were analyzed by the neighbor-joining method and mfa and pra results were divided into 3 groups.

Two mitochondria inheritance-related genes, *rga2* and *rga3*, encoding a mitochondrial targeting signal (MTS), were detected among *U. esculenta* populations, and were located on the a2 and a3 gene complex, respectively (Figure 4). The protein similarity of the 2 Rga proteins was 49%, and that of homologous genes in another smut fungi *Ustanciosporium gigantosporum* was 90%. Rga*2* of *U. esculenta* was much closer to those of other species, including *Sporisorium walker*, *Macalpinomyces eriachnes, U. maydis* and *S. reilianum*, than its own Rga3 protein. As well, rga3 protein of *U. esculenta* showed 66% similarity to *U. xerochloae* protein and 52% similarity to *U. gigantosporum* protein. The 2 *rga* genes might have evolved from 2 different ancestors (Figure 7. However, in another Chinese isolate, MMT, rga3 is missing and is replaced by a transposon-related region (Ye *et al*. 2017). This finding indicates the high activation of transposable elements in the *U. esculenta* population.

**Figure 7.**
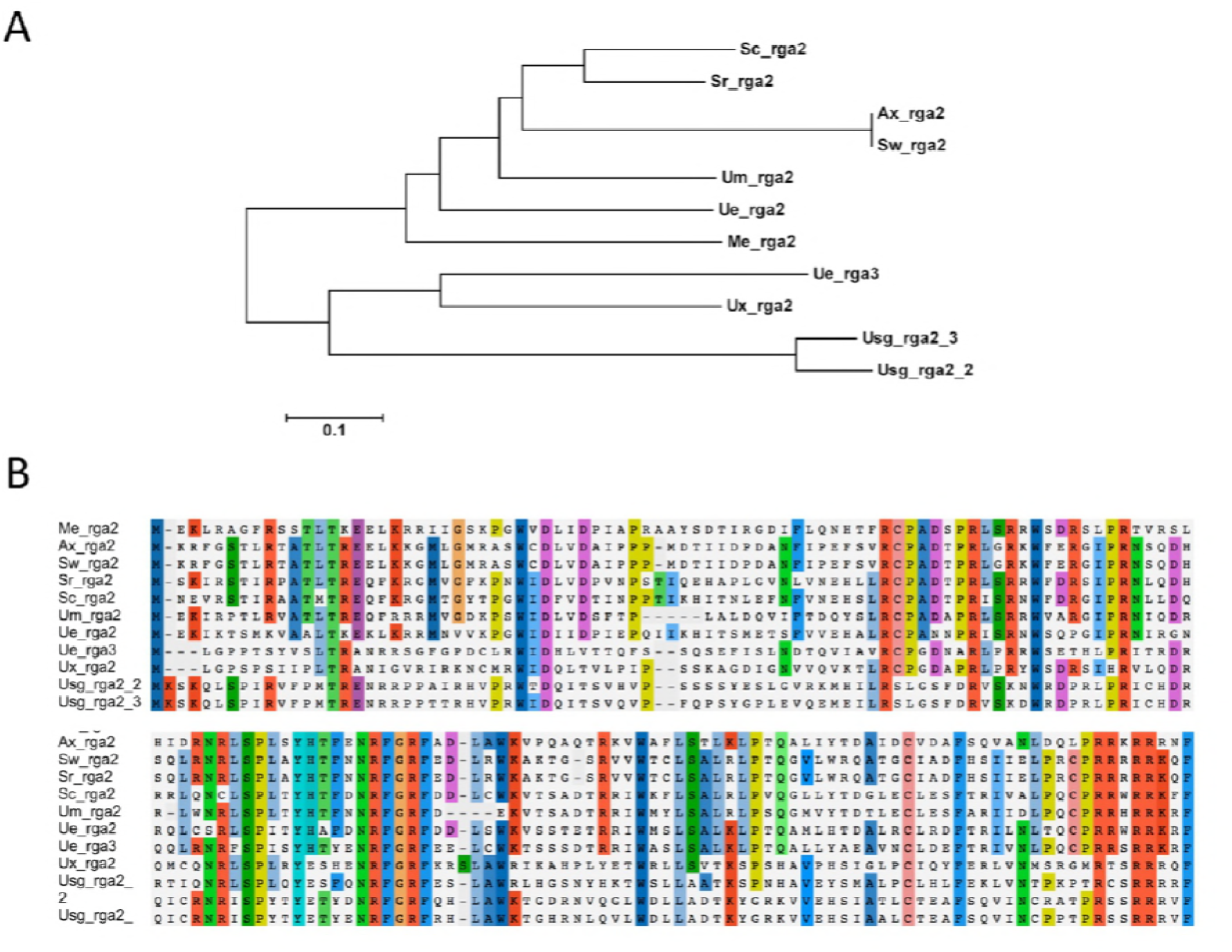
Comparison of rga protein. (A) The phylogenetic tree of rga protein. Two Ue rga proteins were placed in 2 different clades. (B) Comparison of rga homology proteins: Ax, *Anthracocystis walker*; Me, *Macalpinomyces eriachnes*; Sc, *Sporisorium scitamineum;* Ux, *Ustilago xerochloae*; Usg, *Ustanciosporium gigantosporum*; Sw, *Sporisorium walker*. The phylogenetic analysis involved the neighbor-joining method, and the protein matrix for alignment was PAM.

Another genetic variation event occurred on the *MAT* locus. Fot1 family DNA transposon was inserted in the b gene complex. This characteristic was reported in other isolates and was similar to *S. reilianum* SRZ2 and *S. scitamineum* SscI8 but with the opposite site of the b gene complex (Figure 5). Transposons in *U. esculenta* b1 and b2 gene complexes were both 2256 bp and showed 99% similarity. However, the transposon on the b3 gene complex was an exception — 9087 bp — and inserted by several short repeats and sequence. The repeats range from 30 to 69 bp (Figure 5).

### Mating type system of *U. esculenta* is bipolar

Complete *MAT-1* and *MAT-2* loci were retrieved from 12JK1RB1-A1 (*MAT-1*) and UE_mtsf (*MAT-2*) and underwent single molecule real time (SMRT) sequencing. The mating type locus (*MAT*) was identified by using previous identified a and b gene complexes. The *MAT-1* region was covered by 81.15 sequencing reads, on average, which indicated high confidence of its correctness (Figure S1). The sequences within *MAT-1* were variable. *MAT-1* was 555,862 bp and included 115 genes (Table S5) and more than 20 transposable elements predicted by TransposonPSI. About 44.28% featured repeats, with greater proportion than in the non-MAT region (33.13%).

Sequence-region comparison revealed similarity of 2 sequences. *U. esculenta MAT-1* and *MAT-2* sequences showed 66.1% identify by pairwise global alignment (blastn). However, about 61.8% (343,440/555,862 bp) of the *MAT-1* sequence showed synteny to *MAT-2* and more than 95% identity to each other. However, most of the rest of the *MAT-1* region (39.2%) was an intergenic region and occupied by repeats or transposable elements. Some repeats on *MAT-1* were similar (80% to 94% identity) to those of *MAT-2* and spotted on several locations. These sequence phenomena were similar to those for *U. hordei MAT-1*.

Both *U. esculenta* and *U. hordei* mating systems were bipolar but with extremely different mating type locus (*MAT*). Only a partial sequence of *U. esculenta MAT-1* was similar to *U. hordei* chromosome 2. In contrast, *U. esculenta* shared more consensus regions of *MAT-1* and sex chromosome with *U. maydis* (Figure 8) and had a continuous similar sequence with *S. reilianum* (Figure 9). This observation suggested the high relatedness among *U. esculenta*, *U. maydis* and *S. reilianum* during species evolution. Both *U. esculenta* and *U. hordei* underwent a certain genetic event, probably similar, for repeats to have a high portion of mating type locus and to prevent *MAT* loci recombination, so the mating systems are bipolar rather than tetrapolar.

**Figure 8.**
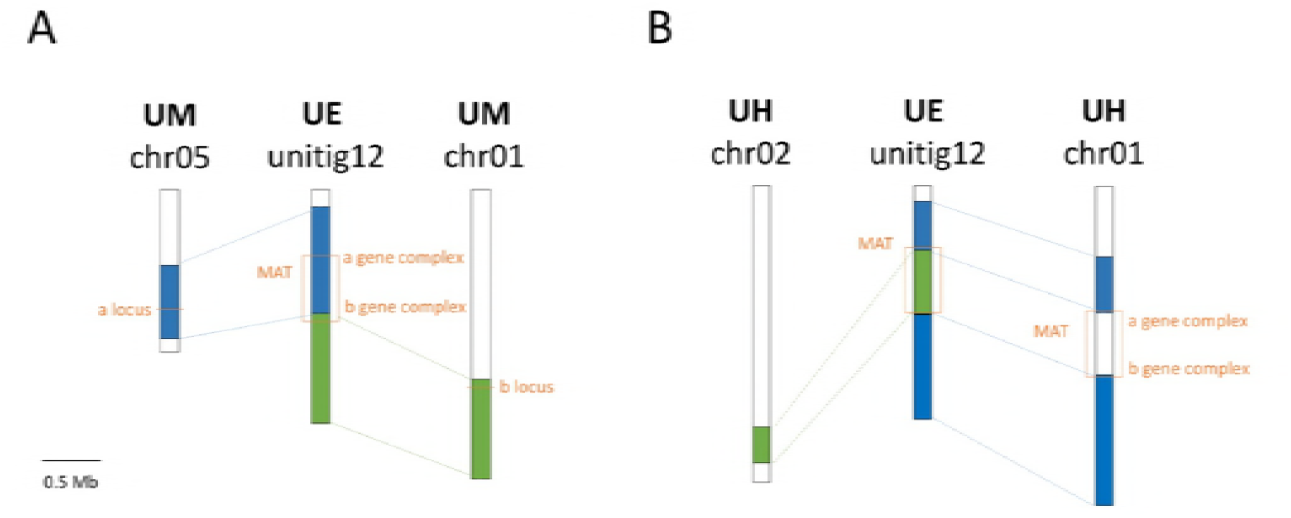
Syntenic comparison of mating-type chromosomes among *U. esculenta, U. maydis* and *U. hordei*. (A) Partial sequence of MAT locus in *U. esculenta* (UE) is syntenic with *U. maydis* chromosomes 5 and 1. (B) Comparison of the sequence of MAT locus between *U. esculenta* and *U. hordei* indicates that these two regions are not similar. *U. esculenta* MAT locus is more similar to that of *U. maydis* than chromosome 2 in *U. hordei*.

**Figure 9.**
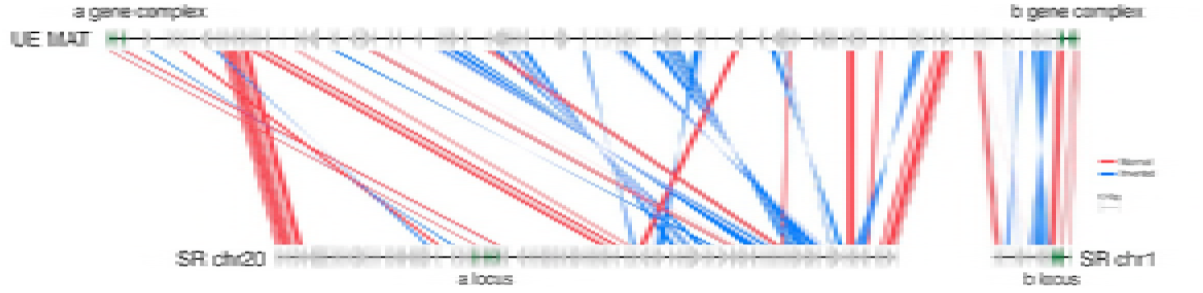
Sequence comparison of *U. esculenta* mating type locus and corresponding region in *S. reilianum*. Green arrows represent mating type-related genes. Red and blue shades are aligned regions where the sequence identity was greater than 70%. Blue indicates inverted sequence. Alignment was by NCBI-blastn with default settings.

### Recombination of mating type locus was suppressed in sexual progenies

*U. maydis* was tetrapolar because the mating type sequence on its four sporidia (progenies) underwent recombination event during meiosis (Kües *et al*. 2011). To discover whether such phenomena occurred on the *MAT* locus of *U. esculenta*, we gathered teliospores from 2 inoculated plants and incubated their sexual progenies to analyze their mating types. The mating analysis involved using complex PCR (Figure 10).

**Figure 10.**
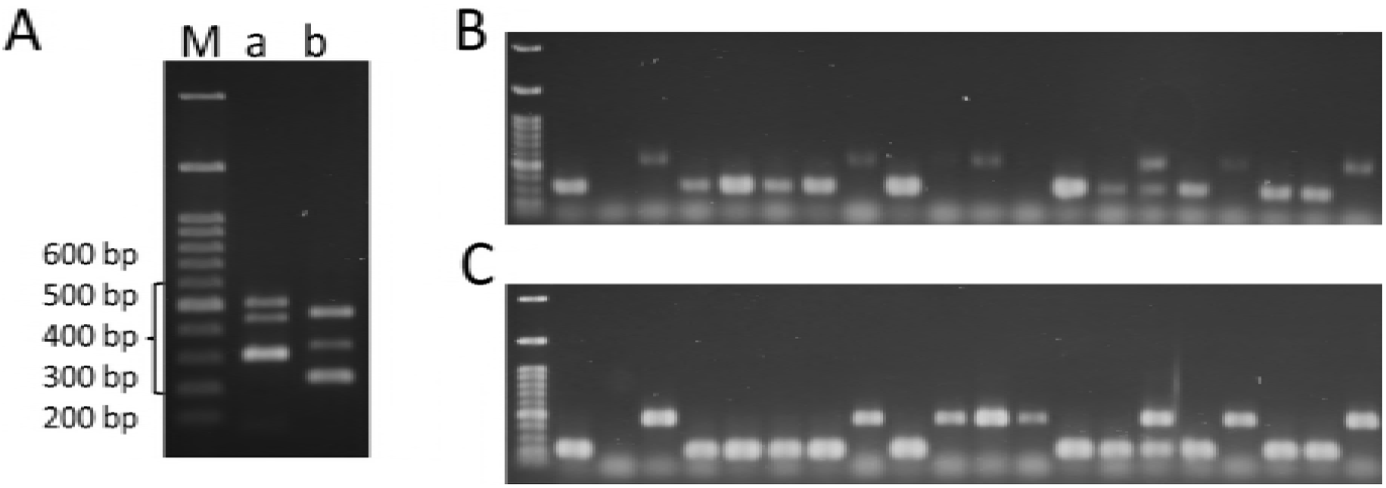
Mating-type screening of *U. esculenta* by multiplex PCR. (A) Three isolates with different mating types were selected to validate the primers for a1/a2/a3 and b1/b2/b3. M: 100 bp marker; Lane 1: PCR products of a1/a2/a3. The amplified fragments were designed in the *pra* genes and the sizes of amplicons were 457 bp for a1, 315 bp for a2 and 515 bp for a3. Lane 2: PCR products of b1/b2/b3. The amplified fragments were designed in the *bE* genes and the sizes of amplicons are 347 bp for b1, 237 bp for b2 and 477 bp for b3. (B) Multiplex PCR screening for mating type showed a2 and a3 mating types among 20 *U. esculenta* isolates. (C) Multiplex PCR screening of the same 20 strains for b mating type showed b2 and b3 mating type and no recombination.

The first group including 466 strains was the offspring of 13PS2GB1-A3 (*MAT-3*) and 13PS2GB1-A4 (*MAT-2*). By using multiplex RCR, we found only 2 mating types, *MAT-2* (a2/b2) and *MAT-3* (a3/b3), which were the same type as their parental strains. Similar results were found in another group [13PJ3GB1-A2 (*MAT-1) ×* 13PJ3GB1-A4 (*MAT-3)]* (Table 2). Because no recombination occurred on the mating type locus during meiosis, the mating system of *U. esculenta* was bipolar.

**Table 2.**
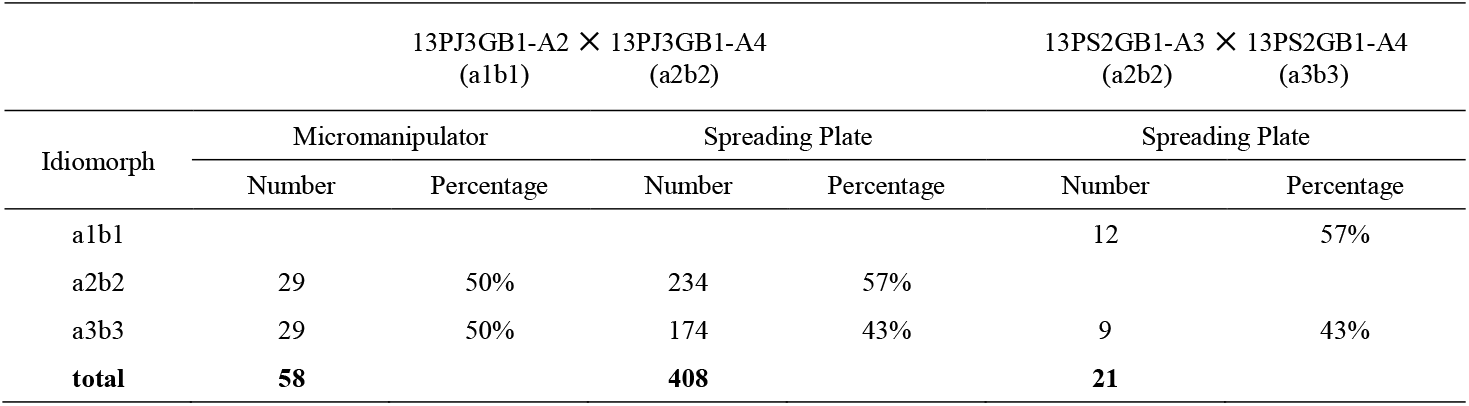
Mating type screening of sexual progenies from the teliospores of two independent inoculated plants

## DISCUSSION

Water bamboo cultivars have been used for long-term artificial selection. However, we do not thoroughly understand the mating system and the impact on gall formation. By studying recombination on the mating type locus and by whole-genome sequencing, we revealed that the mating system of *U. esculenta* is bipolar and has three different mating type loci: *MAT-1* (555,863 bp), *MAT-2* (508,427 bp) and *MAT-3*. The characteristics of *MAT-1* are similar to *U. hordei MAT-1*, including accumulation of insertion elements and different sizes of idiomorphs. Most of *MAT-1* is highly syntenic to sex chromosomes of *U. maydis* and *S. reilianum*, which indicates the occurrence of a recombination event. Several transposable elements (TEs) occur within the region of mating-related genes (a/b gene complexes): one isolate carries *rga*, but the corresponding region on another isolate is replaced by TEs. The other example is the appearance of fot1 family transposons on each B-gene complex, which are located at different positions, as compared with *S. reilianum*.

### *U. esculenta* strain study in Taiwan

In the 145 Taiwanese isolates, three mating types are distributed randomly around Taiwan and can invade both green and red cultivars (Table 3). These observations suggest no distributional differences of mating type in Taiwan.

All collected isolates are from Heishin (T strain), Haushin (unknown strain) or Baishin (MT strain) (Yang and Leu 1978). T strains produce teliospores, but M-T strains do not. With our observations and literature studies, Baishin is prone to transform to Heishin plants under poor weather or in older plants. As compared with Heishin, Baishin is favored by the farmer because of its financial value in the market. However, in this study, we used Heishin isolates for several reasons. Heishin isolates have a complete life cycle and produce teliospores. Its haploid strains are easier to isolate and to use for *in vitro* studies than are Baishin isolates. Furthermore, because it has a natural instinct for producing sexual progeny to finish its life cycle, the M-T strain (Baishin) may be a mutant of the T strain (Heishin).

### Two teliospore-related genes, *hdal* and *rum1*, are slightly different between mtsf and JSKK29

Teliospores form after undergoing a series of filament morphological changes: branching, collapsing, swelling, fragmentation, and teliospore formation (Banuett and Herskowitz 1996). During the procedure, several proteins, Fuz1, Hgl1, (histone deacetylase) Hda1 and Rum1 (homology of human retinoblastoma binding protein 2), are involved in teliospore formation. Gene mutants would cause the absence of teliospores (Chew *et al*. 2008). In our morphological and staining studies of JSKK29, we observed only hyphae branching and partial empty septa. Protein sequence alignment revealed a difference of three peptides between mtsf and JSKK29 in hda1 and rum1, which showed 99.49% and 99.87% identity, respectively. Clarifying the cause of teliospores in the water bamboo needs further study.

### Mating type-related genes reveal the evolutionary information of *U. esculenta*

*U. esculenta* has 3 idiomorphs, *MAT-1, MAT-2* and *MAT-3*. Its gene structure is similar to that of *S. reilianum*, which harbors 2 pheromones on each A mating type locus. We examined all pheromones of *U. esculenta* and related smut fungi (Figure 5). Two pheromones received by the same receptor, such as Mfa3.1 and Mfa2.1 activating Pra1, are very similar regardless of pre- or pro-mature peptides. However, pre-mature peptides of Mfa2.3 and Mfa1.3 at the N-terminal in *U. esculenta* differ. The Mfa2.3 sequence is similar to that of *S. reiliuanum* Mfa1.2 and Mfa3.2, whereas the Mfa1.3 sequence is much closer to that of *S. reiliuanum* Mfa1.3 and Mfa2.3. This divergent sequencing is evidence of the evolutionary remnant between 2 groups of pheromones.

*lga* and *rga* are 2 uniparental mitochondrial-related genes on the A gene complex. Because mutation accumulation usually occurs in mitochondrial inheritance, investigating these 2 genes may reveal clues about the original species of smut fungi (Hoekstra 2000; Fedler *et al*. 2009). Both *U. maydis* and *S. reilianum* carry one *lga* and *rga* on the a2 gene complex, whereas *U. esculenta* harbors one *lga*-like gene and 2 *rga* genes on *MAT-2* and *MAT-3*. The proteins Lga and Rga interfere with mitochondria fusion and regulate pathogenicity in the presence or absence of *mrb1* and *dnm1* (Bortfeld *et al*. 2004; Mahlert *et al*. 2009; Fedler *et al*. 2009). Because *U. esculenta* has the homologous genes of *mrb1* and *dnm1* and its *lga* is not complete as for other smut fungi, *lga* might not be required in *U. esculenta*, which underwent long-term human selection. *U. esculenta* has 2 *rga* genes (*rga2* and *rga3*) on *MAT-2* and *MAT-3. U. gigantosporum* carries 2 Rga proteins as well. However, its genes show high identity (90% identity), whereas those in *U. esculenta* do not (49% identity). As well, *rga3* in isolates from China were lost and replaced by TEs. Because China has a longer cultivar history of water bamboo than Taiwan, this gene variation event indicates that *rga* is less important than other mating-related genes.

### Pheromone precursors of pra3 in *U. esculenta* support that the third R/P system is a variation of other two

*U. esculenta* has 3 pheromone precursors and a pheromone receptor (P/R) system similar to *S. reilianum*. The third pheromone–pheromone receptor system is found only in the basidiomycetes (Kües *et al*. 2011) and is divergent from a common ancestor of other 2 systems. Our phylogenetic analysis (Figure 5B, C) revealed the same result of three pheromone–pheromone receptor systems separated into 3 clades. As well, the pheromone precursors of *U. esculenta* Mfa1.3 and Mfa2.3 differ at the N-terminus. The former shows a closer relationship to other pheromones identified by the Pra2 protein, whereas another pheromone is more similar to Mfa1.3 and Mfa2.3 in *S. reilianum*. This observation supports that the third pheromone–pheromone receptor system is a variation of the other two.

### The mating type locus of *U. esculenta* provides evidence to study the history of *MAT* evolution

*U. esculenta* has a multiple-factor bipolar mating system. Its mating type locus (*MAT*) has several characteristics described in the *U. hordei MAT*, such as accumulation of a repetitive sequence that suppresses the recombination and the size variation of *MAT*, which is caused by the insertion of TEs. The *U. hordei* bipolar system may be created by recombination of the *S. reilianum* (tetrapolar) sex chromosome (Bakkeren *et al*. 2006). However, whether the tetrapolar system evolved from the bipolar system or to the bipolar system is debated (Kües *et al*. 2011).

*MAT-1* and *MAT-2* of *U. esculenta* are 555,863 and 508,427 bp. *MAT-1* is located on the unitig_12, with length of 2,000,476 bp, with one complete telomere. In terms of sequence comparison, a sequence recombination event on large fragments occurred (Figure 8). About half of the *U. esculenta* unitig_12 length is syntenic to *U. maydis* chromosome 1 and the other to chromosome 5. Because the event and outcome are similar to that in *U. hordei*, believed to be degenerated from *S. reilianum*, the sequence event of *U. esculenta* might be the same as *U. hordei:* both species underwent sequence recombination on the sex chromosome. Sex chromosomes of *U. esculenta* and *U. hordei* are very similar, except for the *MAT* regions. Part of the sequence of *U. esculenta MAT* is similar to *U. hordei* chromosome 2, which is in partial synteny to *S. reilianum* chromosome 20. Genes on this syntenic region in *U. hordei* chromosome 2 are dense but are scattered in *U. esculenta* and inserted with repetitive sequences. Some repetitive sequences are shared between these 2 species and located on *MAT* and also all other sequences on the unitig_12 (Figure S2). Comparative analysis revealed that *U. esculenta* underwent a sequence variation event as compared with 3 other smut fungi (*U. maydis, U. hordei, S. reilianum*) and their specialization is closely related. The discovery of the mating type of *U. esculenta* provides evidence to study the evolution of the mating type system.

Many studies have investigated how *U. esculenta* and its host *Z. latifolia* cooperate. The mating type locus is highly related to the development of filaments and controls the process of infection. However, we still do not understand how *U. esculenta* invades its host, how the process of tumor-like tissue formation is related to infection, and whether different mating type conjugations interfere with the size of tumor-like tissue. Such issues are worth of further study.

## Supplemental data

**Figure S1.**
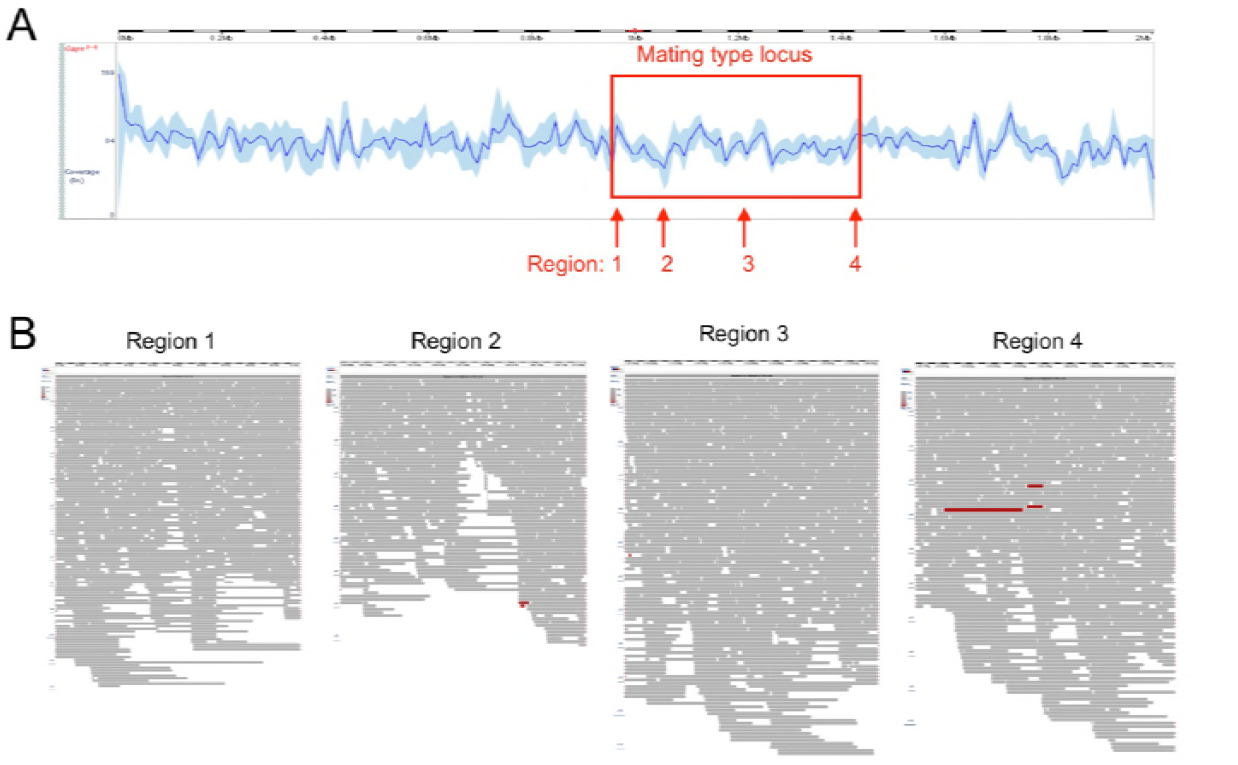
Pacbio reads mapping and assembly result. (A) The average read coverage of unitig_12 where the mating type locus was located. The mean coverage of unitig_12 was 81.15X. The coverage of the MAT locus was similar to other bases of unitig_12. (B) Detailed read-mapping views of mating type locus. Region 1 to 4 represents the regions of start, lowest coverage, and middle and end site, respectively.

**Figure S2.**
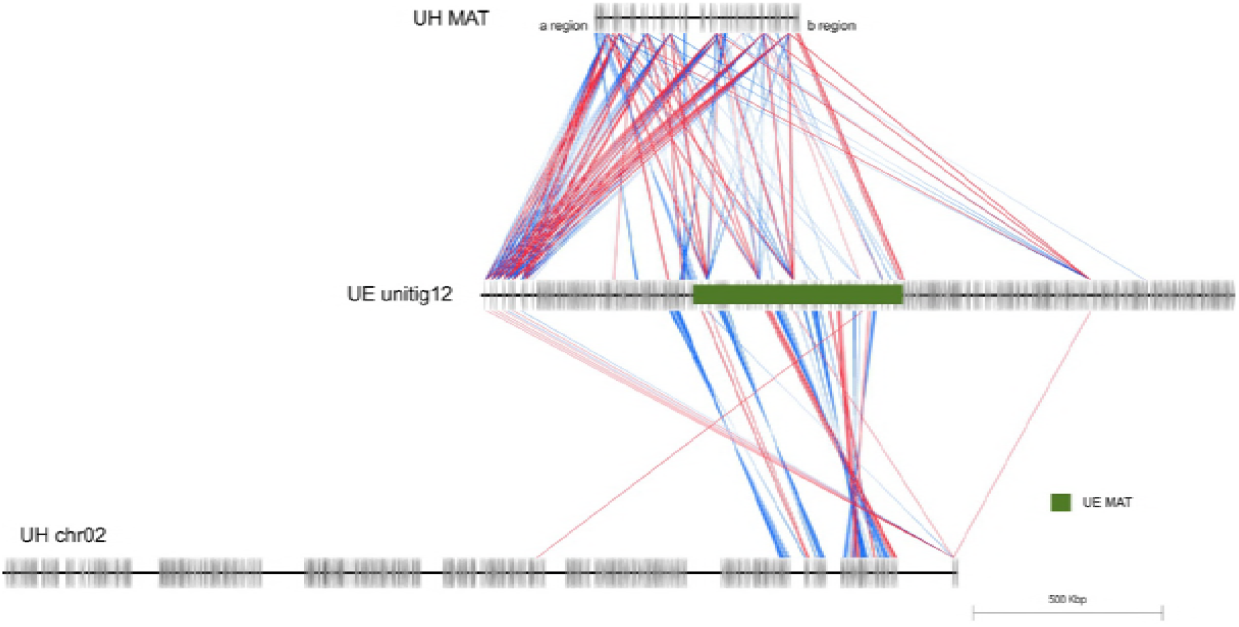
Sequence comparison of *U. esculenta* (UE) MAT on unitig_12 and *U. hordei* (UH) MAT. Green box represents *U. esculenta* MAT with parts of the sequence showing synteny to UH chromosome 2. This region in UH is dense but is scattered in UE. The quantity of the repetitive sequence is shared between sex chromosomes.

**Table S1.** Collections of water bamboo in Taiwan

| Location | Cultivar | Stem type | Collection |
| --- | --- | --- | --- |
| Sanzhi | red shell | heishin | 12SB1RB |
| Sanzhi | red shell | huashin | 12SB1RF |
| Sanzhi | red shell | heishin | 12SF1RB |
| Sanzhi | red shell | heishin | 12SP1RB |
| Sanzhi | red shell | heishin | 12ST1RB |
| Sanzhi | red shell | heishin | 13S1RB |
| Sanzhi | red shell | huashin | 13S1RF |
| Sanzhi | red shell | huashin | 13S2RF |
| Sanzhi | red shell | huashin | 13S3RF |
| Sanzhi | red shell | heishin | 13SS1REB |
| Sanzhi | red shell | huashin | 13SS1REF |
| Sanzhi | red shell | Baishin | 13SS1REW |
| Sanzhi | red shell | heishin | 13SS2RIB |
| Sanzhi | red shell | huashin | 13SS2RIF |
| Sanzhi | red shell | heishin | 13SS3RLB |
| Sanzhi | red shell | huashin | 13SS3RLF |
| Sanzhi | green shell | heishin | 13SS4GEB |
| Sanzhi | green shell | huashin | 13SS4GEF |
| Sanzhi | green shell | heishin | 13SS5GLB |
| Sanzhi | green shell | huashin | 13SS5GLF |
| Jinshan | red shell | heishin | 12JK1RB |
| Jinshan | red shell | heishin | 12JM1RB |
| Jinshan | red shell | huashin | 12JY1RF |
| Jinshan | red shell | heishin | 12JW1RB |
| Jinshan | red shell | heishin | 12JT1RB |
| Taoyuan | Taoyuan No.1 (red shell) | huashin | 13TO1R1F |
| Taoyuan | Taoyuan No.1 (red shell) | heishin | 13TN1R1B |
| Taoyuan | Taoyuan No.1 (red shell) | heishin | 13TD1R1B |
| Taoyuan | Taoyuan No.2 (red shell) | heishin | 13TO1R2B |
| Taoyuan | Taoyuan No.2 (red shell) | heishin | 13TN1R2B |
| Taoyuan | Taoyuan No.2 (red shell) | heishin | 13TD1R2B |
| Puli | green shell | heishin | 13PJ1GB |
| Puli | green shell | heishin | 13PJ2GB |
| Puli | green shell | heishin | 13PJ3GB |
| Puli | green shell | heishin | 13PJ4GB |
| Puli | green shell | heishin | 13PJ5GB |
| Puli | green shell | heishin | 13PJ6GB |
| Puli | green shell | heishin | 13PS1GB |
| Puli | green shell | heishin | 13PS2GB |
| Puli | green shell | heishin | 13PS3GB |

**Table S2.** Strains isolated from germinated teliospores of *U. esculenta*

| Location | Cultivar | Collection | Stem type | Isolate (mating type) |  |  |  |  |  |  |  |
| --- | --- | --- | --- | --- | --- | --- | --- | --- | --- | --- | --- |
| Sanzhi | red shell | 12SB1RB | Heishin | 12SB1RB1-A1* | (MAT-2) | 12SB1RB1-A2* | (MAT-2) | 12SB1RB1-A3* | (MAT-1) | 12SB1RB1-A4* | (MAT-1) |
|  |  |  |  | 12SB1RB1-B1 | (MAT-2) | 12SB1RB1-B2 | (MAT-1) | 12SB1RB1-B3 | (MAT-1) | 12SB1RB1-B4 | (MAT-2) |
|  | early red shell | 13SS1REB1 | Heishin | 13SS1REB1-B1 | (MAT-2) | 13SS1REB1-B2 | (MAT-2) | 13SS1REB1-B3 | (MAT-2) | 13SS1REB1-B4 | (MAT-1) |
|  |  | 13SS2REB2 |  | 13SS1REB2-A1 | (MAT-1) | 13SS1REB2-A3 | (MAT-2) | 13SS1REB2-B1 | (MAT-2) | 13SS1REB2-B2 | (MAT-1) |
|  | green shell | 13SS4GEB | Heishin | 13SS4GEB1-A2 | (MAT-2) | 13SS4GEB1-A3 | (MAT-2) | 13SS4GEB1-B4 | (MAT-1) |  |  |
| Jinshan | red shell | 12JK1RB | Heishin | 12JK1RB1-A1 | (MAT-1) | 12JK1RB1-A2 | (MAT-3) | 12JK1RB1-A3 | (MAT-3) | 12JK1RB1-A4 | (MAT-1) |
| Puli | green shell | 13PJ1GB | Heishin | 13PJ1GB1-A1 | (MAT-2) | 13PJ1GB1-A2 | (MAT-1) | 13PJ1GB1-A3 | (MAT-1) | 13PJ1GB1-A4 | (MAT-2) |
|  |  | 13PJ3GB |  | 13PJ3GB1-A1 | (MAT-3) | 13PJ3GB1-A2 | (MAT-1) | 13PJ3GB1-A3 | (MAT-1) | 13PJ3GB1-A4 | (MAT-3) |
|  |  | 13PS2GB |  | 13PS2GB1-A1 | (MAT-3) | 13PS2GB1-A2 | (MAT-2) | 13PS2GB1-A3 | (MAT-3) | 13PS2GB1-A4 | (MAT-2) |
| Taoyuan | Taoyuan No.1 | 13TN1R1B | Heishin | 13TN1R1B1-A1 | (MAT-3) | 13TN1R1B1-A2 | (MAT-2) | 13TN1R1B1-A3 | (MAT-3) | 13TN1R1B1-A4 | (MAT-2) |
|  |  |  |  | 13TN1R1B1-B1 |  | 13TN1R1B1-B2 |  |  |  |  |  |
|  |  |  |  | 13TN1R1B1-C1 | (MAT-3) | 13TN1R1B1-C2 |  | 13TN1R1B1-C3 | (MAT-3) |  |  |
|  |  |  |  | 13TN1R1B1-D1 | (MAT-3) | 13TN1R1B1-D2 | (MAT-2) | 13TN1R1B1-D3 | (MAT-3) |  |  |
|  | Taoyuan No.2 | 13TO1R2B | Heishin | 13TO1R2B1-A1 | (MAT-1) | 13TO1R2B1-A2 | (MAT-2) | 13TO1R2B1-A3 | (MAT-1) | 13TO1R2B1-A4 | (MAT-2) |
|  |  |  |  | 13TO1R2B1-B1 | (MAT-2) | 13TO1R2B1-B2 | (MAT-1) | 13TO1R2B1-B3 | (MAT-2) | 13TO1R2B1-B4 | (MAT-1) |
|  |  |  |  | 13TO1R2B1-C1 | (MAT-2) | 13TO1R2B1-C2 | (MAT-2) | 13TO1R2B1-C3 | (MAT-2) | 13TO1R2B1-C4 |  |

\*A1, A2, A3, A4 indicate the 4 sporidia isolated from the same teliospore; B1, B2, B3, B4 are derived from the same teliospore.

**Table S3.**
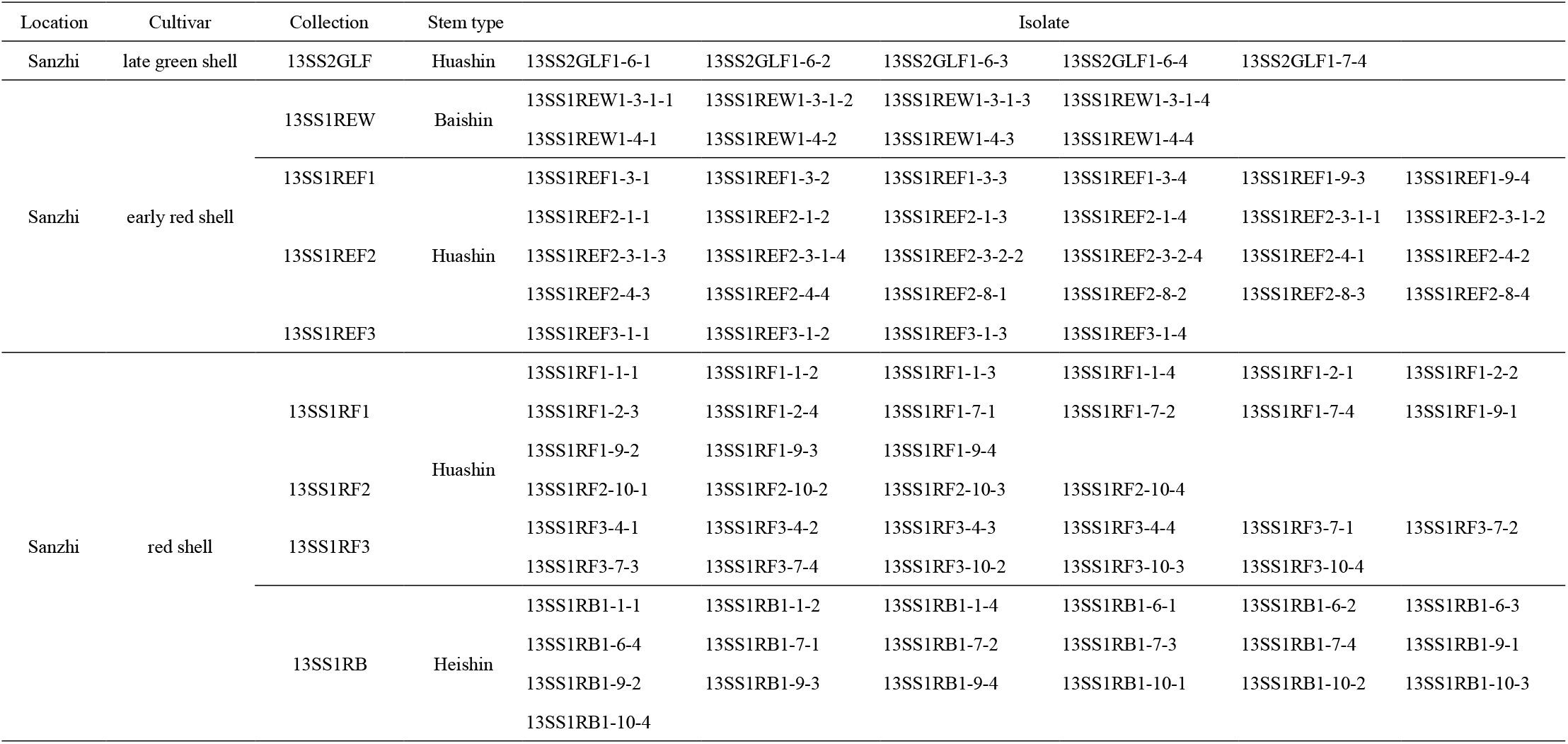
Strains isolated from stem tissue of water bamboo

**Table S4.**
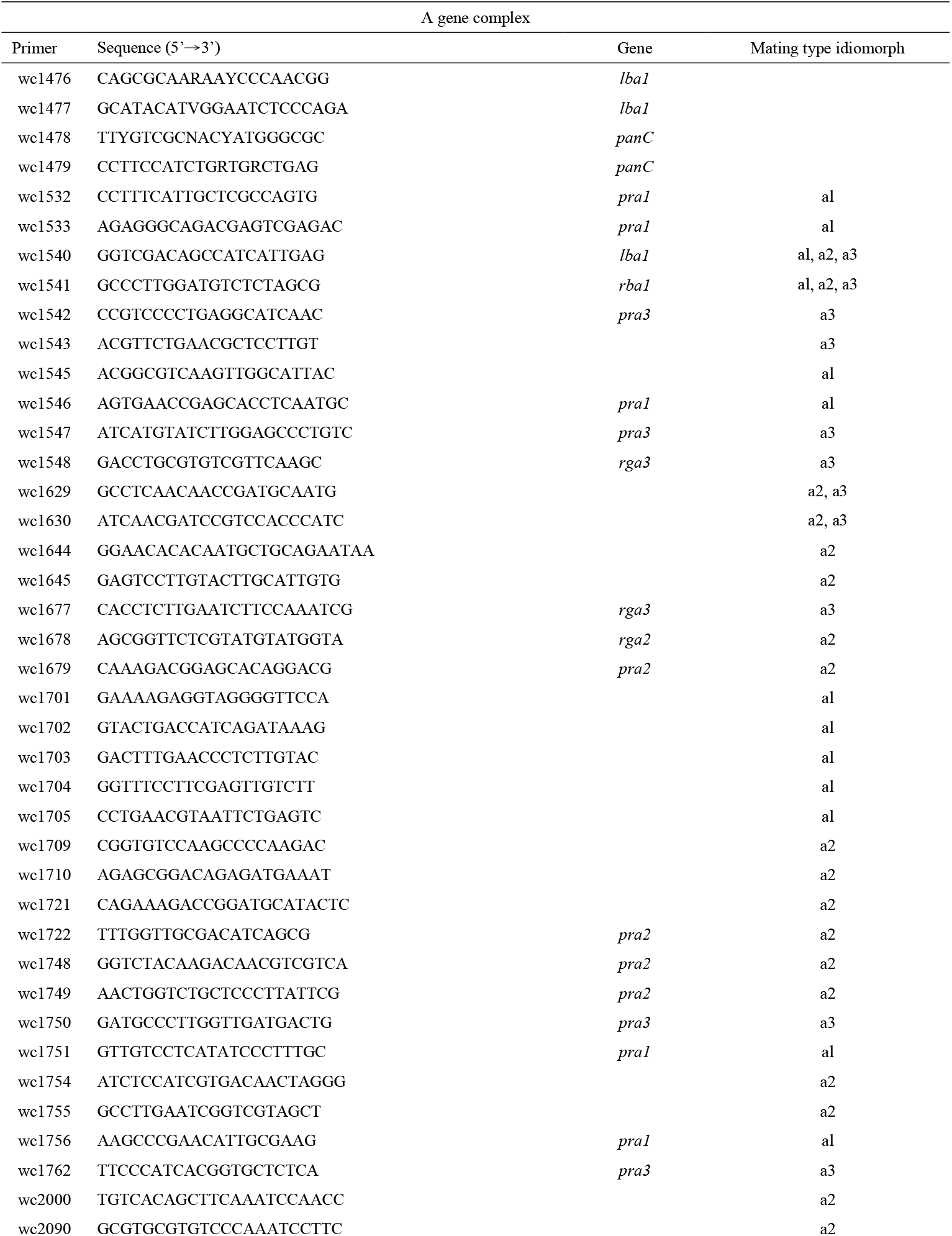
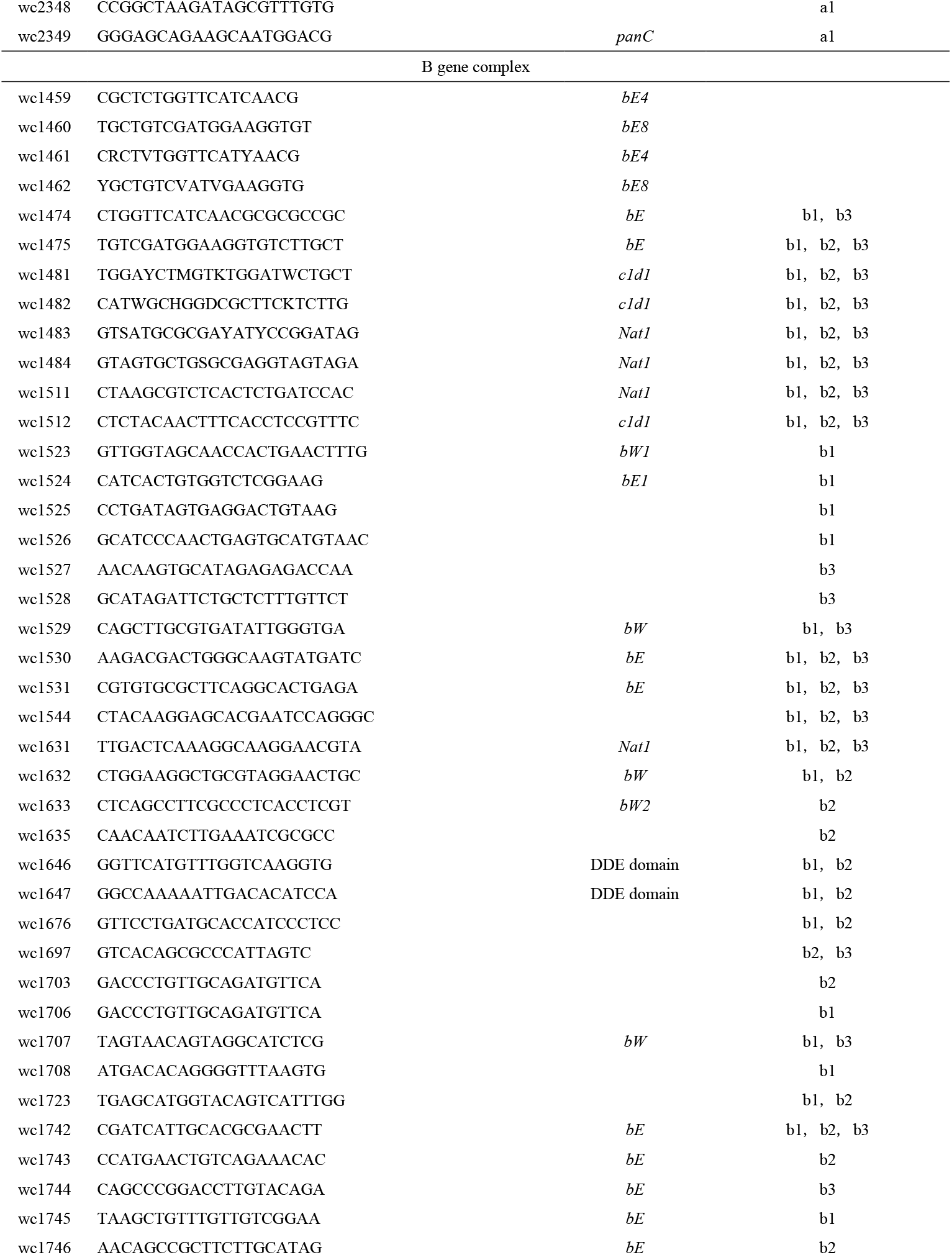
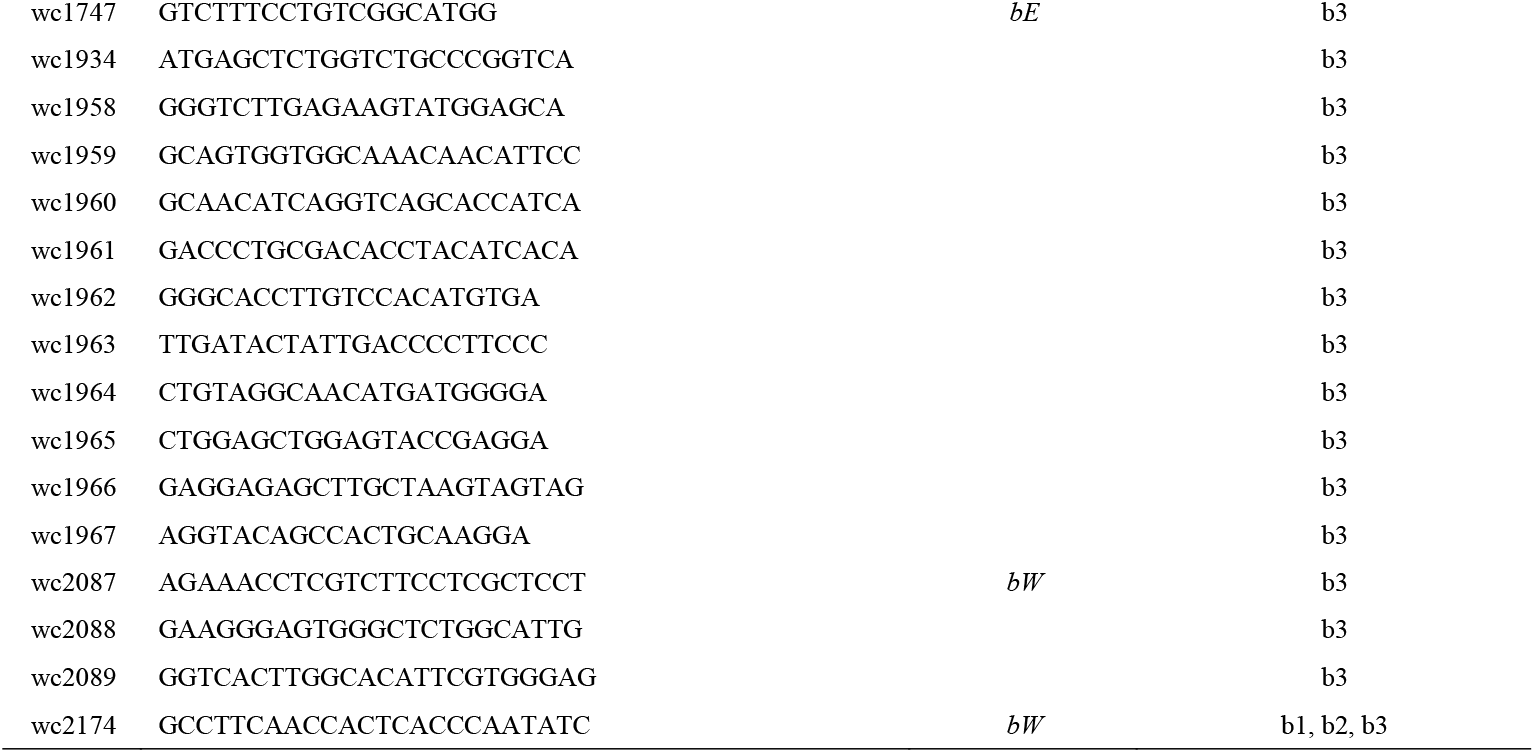
Primer sets of mating type A and B gene complex

**Table S5.**
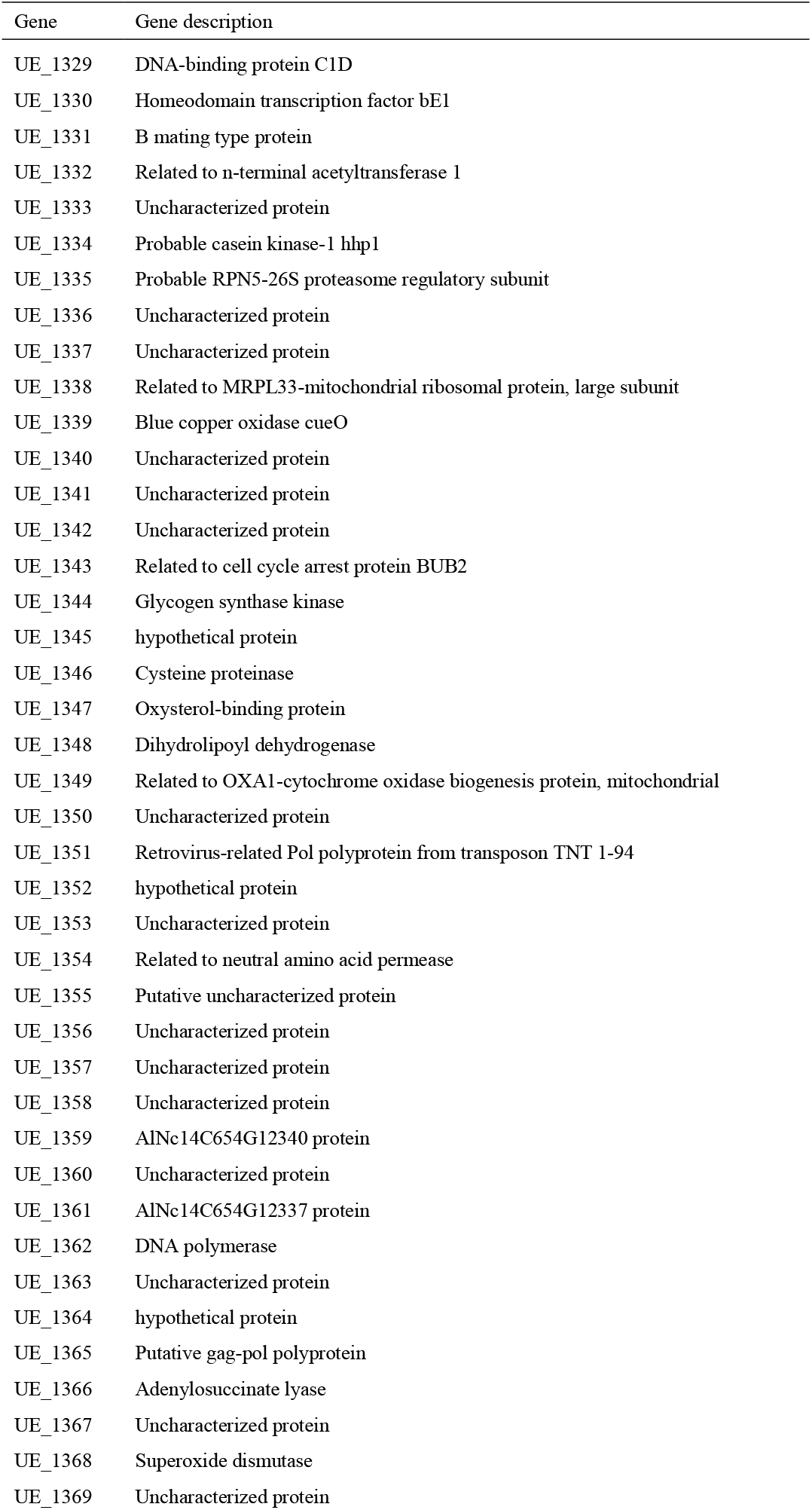
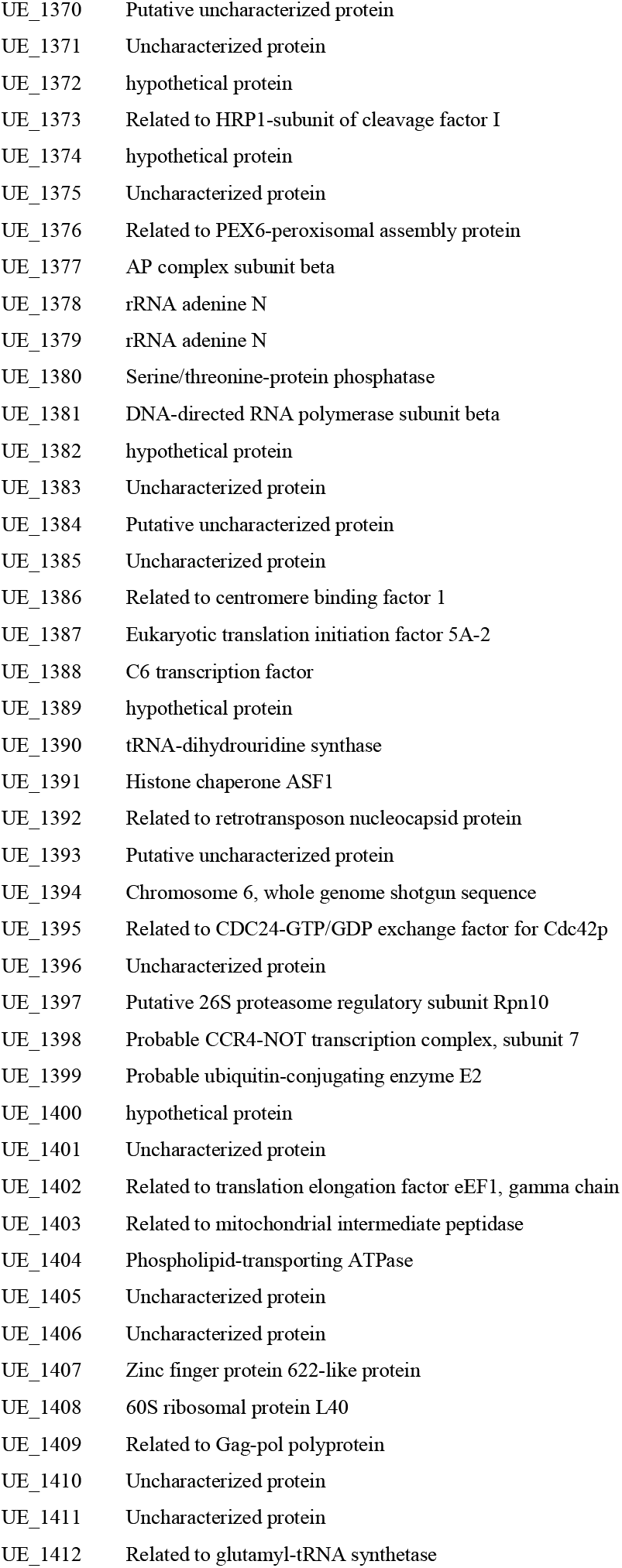
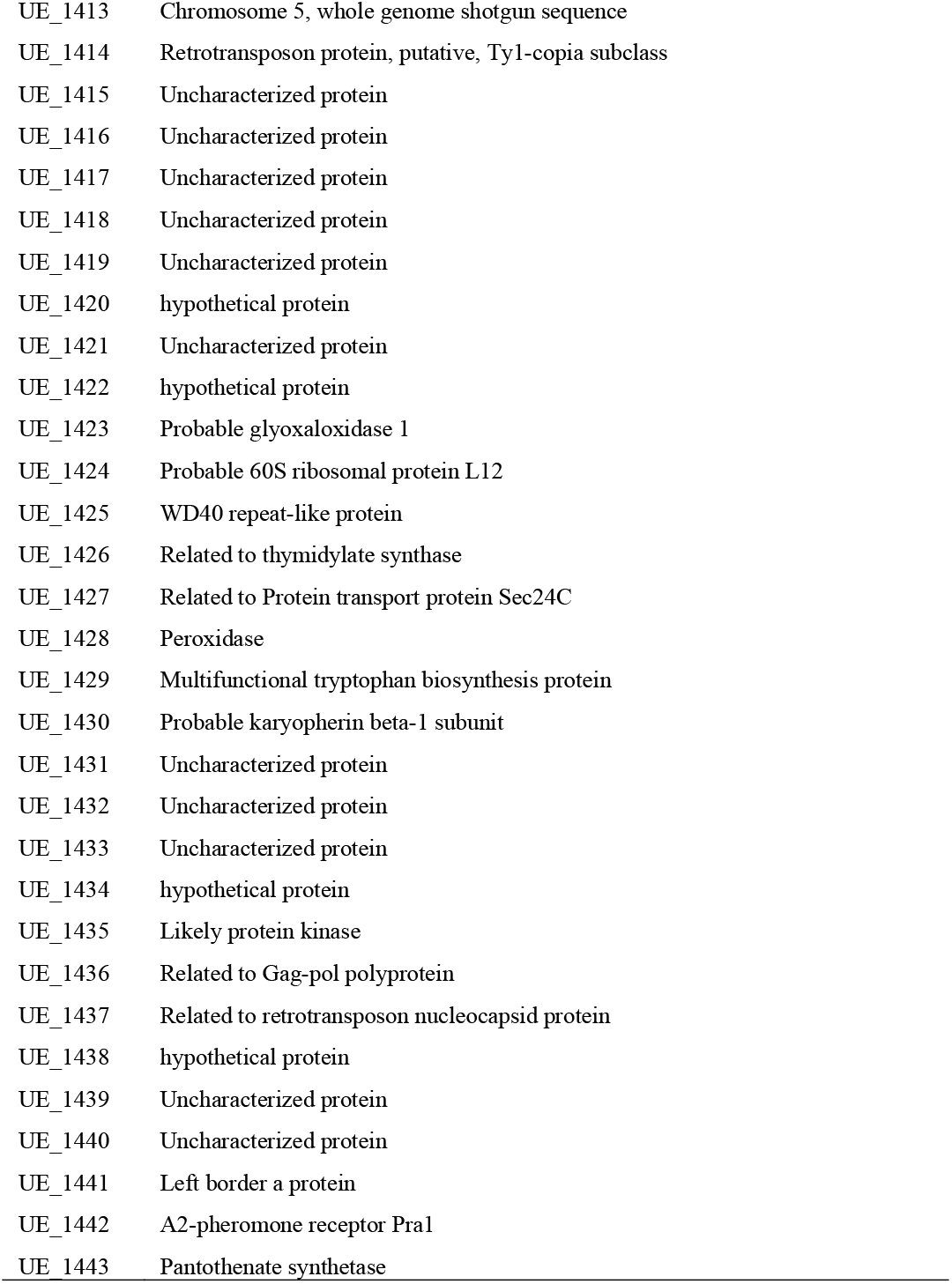
Predicted genes on the *MAT-1* locus

| Gene | Gene description |
| --- | --- |
| UE_1329 | DNA-binding protein C1D |
| UE_1330 | Homeodomain transcription factor bE1 |
| UE_1331 | B mating type protein |
| UE_1332 | Related to n-terminal acetyltransferase 1 |
| UE_1333 | Uncharacterized protein |
| UE_1334 | Probable casein kinase-1 hhp1 |
| UE_1335 | Probable RPN5-26S proteasome regulatory subunit |
| UE_1336 | Uncharacterized protein |
| UE_1337 | Uncharacterized protein |
| UE_1338 | Related to MRPL33-mitochondrial ribosomal protein, large subunit |
| UE_1339 | Blue copper oxidase cueO |
| UE_1340 | Uncharacterized protein |
| UE_1341 | Uncharacterized protein |
| UE_1342 | Uncharacterized protein |
| UE_1343 | Related to cell cycle arrest protein BUB2 |
| UE_1344 | Glycogen synthase kinase |
| UE_1345 | hypothetical protein |
| UE_1346 | Cysteine proteinase |
| UE_1347 | Oxysterol-binding protein |
| UE_1348 | Dihydrolipoyl dehydrogenase |
| UE_1349 | Related to OXA1-cytochrome oxidase biogenesis protein, mitochondrial |
| UE_1350 | Uncharacterized protein |
| UE_1351 | Retrovirus-related Pol polyprotein from transposon TNT 1-94 |
| UE_1352 | hypothetical protein |
| UE_1353 | Uncharacterized protein |
| UE_1354 | Related to neutral amino acid permease |
| UE_1355 | Putative uncharacterized protein |
| UE_1356 | Uncharacterized protein |
| UE_1357 | Uncharacterized protein |
| UE_1358 | Uncharacterized protein |
| UE_1359 | AINc14C654G12340 protein |
| UE_1360 | Uncharacterized protein |
| UE_1361 | AINc14C654G12337 protein |
| UE_1362 | DNA polymerase |
| UE_1363 | Uncharacterized protein |
| UE_1364 | hypothetical protein |
| UE_1365 | Putative gag-pol polyprotein |
| UE_1366 | Adenylosuccinate lyase |
| UE_1367 | Uncharacterized protein |
| UE_1368 | Superoxide dismutase |
| UE_1369 | Uncharacterized protein |
| UE_1370 | Putative uncharacterized protein |
| UE_1371 | Uncharacterized protein |
| UE_1372 | hypothetical protein |
| UE_1373 | Related to HRP1-subunit of cleavage factor I |
| UE_1374 | hypothetical protein |
| UE_1375 | Uncharacterized protein |
| UE_1376 | Related to PEX6-peroxisomal assembly protein |
| UE_1377 | AP complex subunit beta |
| UE_1378 | rRNA adenine N |
| UE_1379 | rRNA adenine N |
| UE_1380 | Serine/threonine-protein phosphatase |
| UE_1381 | DNA-directed RNA polymerase subunit beta |
| UE_1382 | hypothetical protein |
| UE_1383 | Uncharacterized protein |
| UE_1384 | Putative uncharacterized protein |
| UE_1385 | Uncharacterized protein |
| UE_1386 | Related to centromere binding factor 1 |
| UE_1387 | Eukaryotic translation initiation factor 5A-2 |
| UE_1388 | C6 transcription factor |
| UE_1389 | hypothetical protein |
| UE_1390 | tRNA-dihydrouridine synthase |
| UE_1391 | Histone chaperone ASF1 |
| UE_1392 | Related to retrotransposon nucleocapsid protein |
| UE_1393 | Putative uncharacterized protein |
| UE_1394 | Chromosome 6, whole genome shotgun sequence |
| UE_1395 | Related to CDC24-GTP/GDP exchange factor for Cdc42p |
| UE_1396 | Uncharacterized protein |
| UE_1397 | Putative 26S proteasome regulatory subunit Rpn10 |
| UE_1398 | Probable CCR4-NOT transcription complex, subunit 7 |
| UE_1399 | Probable ubiquitin-conjugating enzyme E2 |
| UE_1400 | hypothetical protein |
| UE_1401 | Uncharacterized protein |
| UE_1402 | Related to translation elongation factor eEF1, gamma chain |
| UE_1403 | Related to mitochondrial intermediate peptidase |
| UE_1404 | Phospholipid-transporting ATPase |
| UE_1405 | Uncharacterized protein |
| UE_1406 | Uncharacterized protein |
| UE_1407 | Zinc finger protein 622-like protein |
| UE_1408 | 60S ribosomal protein L40 |
| UE_1409 | Related to Gag-pol polyprotein |
| UE_1410 | Uncharacterized protein |
| UE_1411 | Uncharacterized protein |
| UE_1412 | Related to glutamyl-tRNA synthetase |
| UE_1413 | Chromosome 5, whole genome shotgun sequence |
| UE_1414 | Retrotransposon protein, putative, Ty1-copia subclass |
| UE_1415 | Uncharacterized protein |
| UE_1416 | Uncharacterized protein |
| UE_1417 | Uncharacterized protein |
| UE_1418 | Uncharacterized protein |
| UE_1419 | Uncharacterized protein |
| UE_1420 | hypothetical protein |
| UE_1421 | Uncharacterized protein |
| UE_1422 | hypothetical protein |
| UE_1423 | Probable glyoxaloxidase 1 |
| UE_1424 | Probable 60S ribosomal protein L12 |
| UE_1425 | WD40 repeat-like protein |
| UE_1426 | Related to thymidylate synthase |
| UE_1427 | Related to Protein transport protein Sec24C |
| UE_1428 | Peroxidase |
| UE_1429 | Multifunctional tryptophan biosynthesis protein |
| UE_1430 | Probable karyopherin beta-1 subunit |
| UE_1431 | Uncharacterized protein |
| UE_1432 | Uncharacterized protein |
| UE_1433 | Uncharacterized protein |
| UE_1434 | hypothetical protein |
| UE_1435 | Likely protein kinase |
| UE_1436 | Related to Gag-pol polyprotein |
| UE_1437 | Related to retrotransposon nucleocapsid protein |
| UE_1438 | hypothetical protein |
| UE_1439 | Uncharacterized protein |
| UE_1440 | Uncharacterized protein |
| UE_1441 | Left border a protein |
| UE_1442 | A2-pheromone receptor Pra1 |
| UE_1443 | Pantothenate synthetase |

